# Rewiring of chromatin loops in adipogenesis reveals targets for obesity and diabetes intervention

**DOI:** 10.1101/2023.06.27.546683

**Authors:** Ionel Sandovici, Borbala Mifsud, Katherine A. Kentistou, Amy Emery, Pawan Gulati, Ayesha Banu, Niamh Campbell, Bryn S. Hardwick, Alex T. Crooks, Denise S. Fernandez-Twinn, Lais V. Mennitti, Luma Srour, Sherine Awad, Davide Chiarugi, Russell S. Hamilton, Steven W. Wingett, Peter Fraser, Ken K. Ong, Stefan Schoenfelder, Farhan Mohammad, Stephen O’Rahilly, John R.B. Perry, Ashok R. Venkitaraman, Susan E. Ozanne, Miguel Constância

**Affiliations:** Metabolic Research Laboratory, Wellcome-MRC Institute of Metabolic Science, University of Cambridge School of Clinical Medicine, Cambridge CB2 0QQ, UK; Department of Obstetrics and Gynaecology and National Institute for Health Research Cambridge Biomedical Research Centre, Cambridge, CB2 0SW, UK; Centre for Trophoblast Research, Department of Physiology, Development and Neuroscience, University of Cambridge, Cambridge CB2 3EG, UK; Division of Genomics and Translational Biomedicine, College of Health and Life Sciences, Hamad Bin Khalifa University, Education City, Doha, Qatar; William Harvey Research Institute, Queen Mary University of London, London, UK; MRC Epidemiology Unit, University of Cambridge School of Clinical Medicine, Institute of Metabolic Science, Cambridge, CB2 0QQ, UK; Medical Research Council Cancer Unit, University of Cambridge, Hills Road, Cambridge, CB2 0XZ, UK; Division of Biological & Biomedical Sciences (BBS), College of Health & Life Sciences (CHLS), Hamad Bin Khalifa University (HBKU), Doha, Qatar; Department of Genetics, University of Cambridge, Downing Street, Cambridge, CB2 3EH, UK; Nuclear Dynamics Programme, The Babraham Institute, Cambridge, CB22 3AT, UK; MRC Laboratory of Molecular Biology, Cambridge, UK; Department of Biological Science, Florida State University, Tallahassee, FL, USA; Epigenetics Programme, Babraham Institute, Cambridge CB22 3AT, UK; Cancer Science Institute of Singapore, National University of Singapore, 14 Medical Drive, Singapore 117599, Singapore; Institute for Molecular & Cell Biology, Agency for Science, Technology and Research (A*STAR), 61 Biopolis Drive, Singapore - 138673

## Abstract

Adipogenesis is a multi-stage process essential for healthy fat storage and metabolic regulation. While early regulatory mechanisms are well characterized, the control of late-stage adipocyte differentiation remains poorly understood. Integrating CAGE-seq, promoter capture Hi-C, and a high-throughput siRNA screen of druggable genes, we report here that chromatin architecture rewiring promotes gene regulation changes essential for terminal adipogenesis. We identified nine clusters of dynamic promoter-anchored chromosomal interactions, many involving distal enhancers. Functional screening of genes engaged in these interactions revealed 19 novel regulators of late adipogenesis, including proteins with peptidase and ubiquitin ligase activity. Human genetic variant-to-gene mapping, coupled with cross-species chromatin interaction and synteny analyses, highlighted new gene-trait associations relevant to lipid traits (FXYD5, LAP3, SGPP1) and type 2 diabetes (FBXO17, FN3KRP, ZFAND6, TTC3). Our findings define the 3D gene regulatory landscape of late adipogenesis. The molecular links uncovered here provide mechanistic insight into metabolic disease risk and potential interventions.

## Main

The global rise in obesity has intensified efforts to understand the mechanisms governing adipocyte differentiation and maturation. Adipogenesis is a multistep process beginning with the commitment of mesenchymal precursors to pre-adipocytes, followed by terminal differentiation into insulin-responsive mature adipocytes through cell cycle arrest and sustained lipogenesis^1^. Much of our understanding of adipogenesis comes from in vitro models, including the murine 3T3-L1^2^ and OP9-K^3^ cell lines, which differentiate in response to adipogenic stimuli. In 3T3-L1 cells, adipogenesis proceeds through early (0–36 h), intermediate (36–72 h), and late (days 3–7) stages^4^. Notably, these in vitro stages recapitulate key features of in vivo adipogenesis, as confirmed by lineage tracing studies in mice^5,6^.

While the molecular regulation of early and intermediate stages of adipocyte differentiation is well characterized, little is known about the mechanisms driving the late stages of this process— particularly those involving epigenetic control. This knowledge gap is notable given that adipocyte size, number, and turnover are major determinants of fat mass and strongly influence metabolic health. Large, hypertrophic adipocytes are frequently associated with insulin resistance, dyslipidemia, hypertension, and increased risk of type 2 diabetes^7,8^.

Epigenetic regulation plays a critical role in cell differentiation, especially via enhancers—regulatory DNA elements that modulate gene expression irrespective of their position relative to target promoters^9^. H3K4me1 marks many enhancers regardless of activity, while its combination with H3K27ac or H3K27me3 indicates active or poised enhancers, respectively^9^. Enhancers act on target promoters through physical interactions enabled by 3D chromatin folding^10^. Promoter capture Hi-C (PCHi-C), an adaptation of chromosome conformation capture (3C) technology^11^, allows genome-wide mapping of promoter interactions and has been applied to both mouse^12^ and human^13^ genomes. In 3T3-L1 cells, early and intermediate differentiation stages show rapid and extensive rewiring of promoter-anchored chromatin loops^14^, yet changes during late-stage differentiation remain unexplored.

To address this knowledge gap, we generated novel PCHi-C and CAGE-seq data to examine promoter-anchored chromatin loops and TSS activity, and integrated it with publicly available ChIP-seq and WGBS data to analyse histone marks and DNA methylation during 3T3-L1 adipocyte differentiation. Building on these results, we performed a high-throughput siRNA screen targeting >2,900 druggable genes in OP9-K cells, quantifying lipid accumulation as a readout of adipogenic potential. By integrating these data, we aimed to identify regulators of terminal adipocyte differentiation.

To further assess functional relevance, we validated selected genes in vivo via siRNA knockdown in the Drosophila fat body, and mined mouse phenotyping (MGI, IMPC) and human GWAS data for associations with obesity-related traits. Our analyses uncovered novel regulators of late-stage adipogenesis, including multiple genes with peptidase or ubiquitin ligase activity, offering new insights into adipocyte biology and paving the way for potential novel therapeutic targets.

## Results

### Dynamic promoter usage and chromatin looping shape the adipogenic transcriptional program

We differentiated mouse 3T3-L1 pre-adipocytes^15^ (female; see Methods) for seven days in adipogenic media, resulting in uniform lipid droplet accumulation and altered expression of key adipogenic markers^16,17^, including upregulation of Adipoq, rJCebpa, and rJPpargrJand downregulation ofrJZfp521 (Extended Data Fig. 1a). We profiled transcription start site (TSS) activity at day 0 (D0) and day 7 (D7) using nAnT-iCAGE (CAGE-seq)^18^, identifying over 118,000 TSSs (Supplementary Table 1a). Approximately 20% were differentially expressed (Fig. 1a), corresponding to ∼7,800 differentially expressed genes (DEGs; Supplementary Table 1b), with two-thirds downregulated during differentiation. qRT-PCR of 12 DEGs in independent replicates confirmed strong correlation with CAGE-seq data (Extended Data Fig. 1b).

**Fig. 1:**
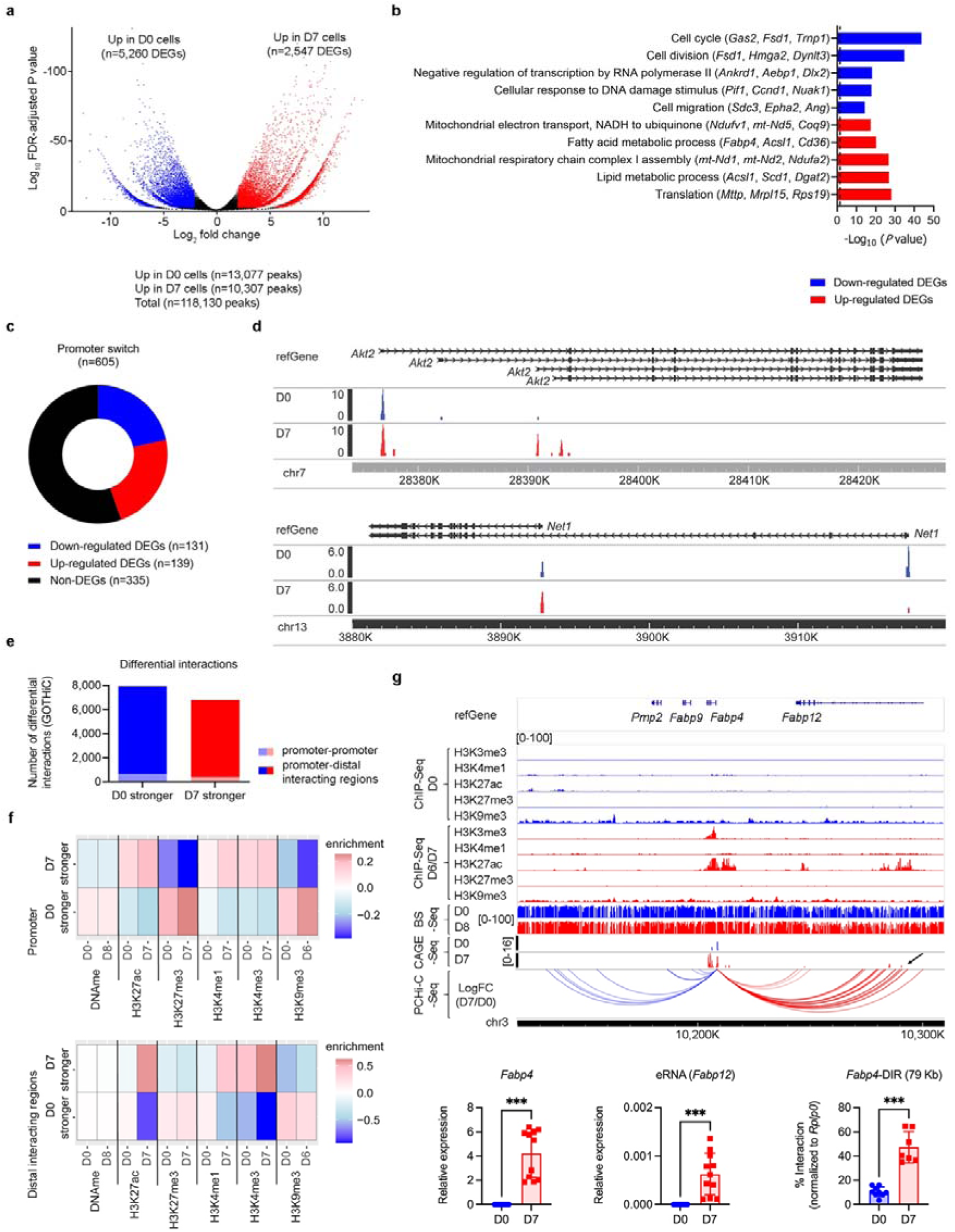
Transcriptional and epigenetic changes during in vitro adipogenesis in the 3T3-L1 model. a, Volcano plot showing differentially expressed transcripts between day 0 (D0) and day 7 (D7) of 3T3-L1 differentiation, as detected by CAGE-seq. A total of 23,300 transcripts (∼16.5%) from 7,807 genes were differentially expressed. b, Gene ontology enrichment analysis (DAVID) for biological processes associated with differentially expressed genes (fold change >2). Dashed line indicates FDR-adjusted P1<10.05. c, Distribution of promoter switching events among 605 genes showing alternative promoter usage during adipogenesis, relative to overall gene expression levels. d, Genome browser examples of two genes undergoing promoter switching, with CAGE-seq peaks expressed at D0 (blue) and D7 (red). e, Classification of D0/D7 differential promoter–anchored chromatin interactions identified by promoter capture Hi-C. The majority of differential loops involve distal intergenic or intragenic regions, with a minority representing promoter–promoter contacts. f, Chromatin marks at promoters (top) and distal interacting regions (bottom) involved in differential D0/D7 loops, showing acquisition of enhancer-associated histone modifications during adipogenesis. g, Top: Example of a D7-specific loop between the Fabp4 promoter and a downstream enhancer-like region located ∼79 kb away within the Fabp12 locus, marked by H3K27ac and eRNA transcription. Red and blue arcs indicate interactions stronger and weaker at D7, respectively. Bottom: qRT-PCR validation of Fabp4 mRNA and eRNA expression and q3C validation of Fabp4–distal interaction region (DIR) contact. Data represent mean ± s.d.; ***P1<10.001 by Mann–Whitney test.

Gene ontology (GO) analysis revealed that downregulated genes were enriched in cell cycle, transcription, DNA repair, and migration, while upregulated genes were associated with lipid metabolism, mitochondrial function, and translation (Fig. 1b; Supplementary Tables 1c–f). Over 600 genes exhibited promoter switching, defined as reciprocal expression changes of multiple TSSs from the same gene during the transition from pre-adipocytes to differentiated adipocytes (Fig. 1c; Supplementary Table 1g). These were enriched in pathways related to transcriptional and chromatin regulation (Extended Data Fig. 1c; Supplementary Tables 1h,i). Examples of genes showing promoter switching during adipogenesis include Akt2 (insulin signalling), Net1 (cell motility and mitosis), Fhl1 (proliferation and signalling), Jmjd1c (histone demethylation), and Smarcd2 (chromatin remodelling) (Fig. 1d; Extended Data Fig. 1d).

To assess in vivo relevance, we isolated GFP^+^ committed pre-adipocytes and mature adipocytes from the gonadal fat of male Zfp423^GFP^ reporter mice^19^ using FACS, and performed RNA-seq. We identified 5,174 DEGs in vivo, of which ∼2,500 overlapped with CAGE-seq DEGs (Supplementary Table 1j). Over 85% of shared DEGs changed in the same direction during adipogenesis, with strong positive correlation (Extended Data Fig. 1e), supporting the utility of the 3T3-L1 model for studying molecular features of in vivo adipogenesis.

To investigate changes in 3D genome architecture, we performed promoter capture Hi-C (PCHi-C)^12^ in D0 and D7 3T3-L1 cells. GOTHiC analysis^20^ identified 14,723 differential interactions (DInts) between single HindIII fragments (Supplementary Table 2a). Over 93% of DInts connected promoters to distal non-promoter regions (Fig. 1e). These changes coincided with shifts in epigenetic landscapes. At promoters, we observed dynamic changes in repressive marks H3K27me3 and H3K9me3, and modest changes in H3K4me1. At distal regions, marks associated with enhancer (H3K27ac, H3K4me1) and transcriptional (H3K4me3) activity were most affected (Fig.1f). We selected six DInts spanning distances from 61 kb to 1.24 Mb for validation using quantitative 3C (q3C) assays^21^. All showed significant interaction changes between D0 and D7 in the same direction as predicted by GOTHiC (Fig. 1g; Extended Data Fig. 1f). Among them, a DInt linking the Fabp4 promoter with an intronic region of Fabp12—previously reported during adipogenesis^14,22^—was significantly strengthened at D7. This intronic region simultaneously gained H3K27ac and evidence of eRNA transcription, consistent with enhancer activation (Fig.1g).

Collectively, these results reveal an unrecognized programme of coordinated transcriptional changes that underlie 3T3-L1 adipogenesis. These coordinated transcriptional changes are mediated by alterations in chromatin architecture, wherein changes in epigenetic marks accompany chromatin interaction remodelling and dynamic promoter usage. Altogether, these findings uncover a previously underappreciated layer of regulatory complexity and point to potential implications for adipose tissue development and metabolic disease.

### Temporal clustering of promoter-anchored loops reveals enhancer engagement during late-stage adipogenesis

We therefore characterized the dynamic promoter-anchored chromatin interactions during adipocyte differentiation by analysing PCHi-C data from 3T3-L1 cells at four timepoints (D0, 4h, D2, D7), integrating our dataset with that from Siersbæk et al.^14^. Using GOTHiC and k-means clustering, we identified 23,606 DInts, grouped into nine clusters with distinct temporal profiles (Fig. 2a; Supplementary Table 2b). Clusters 1, 2, 6, and 8 showed transient interactions in early or intermediate differentiation. Clusters 4 and 9 included DInts that were strengthened during terminal adipogenesis, while clusters 3, 5, and 7 exhibited reduced interactions at later stages.

**Fig. 2:**
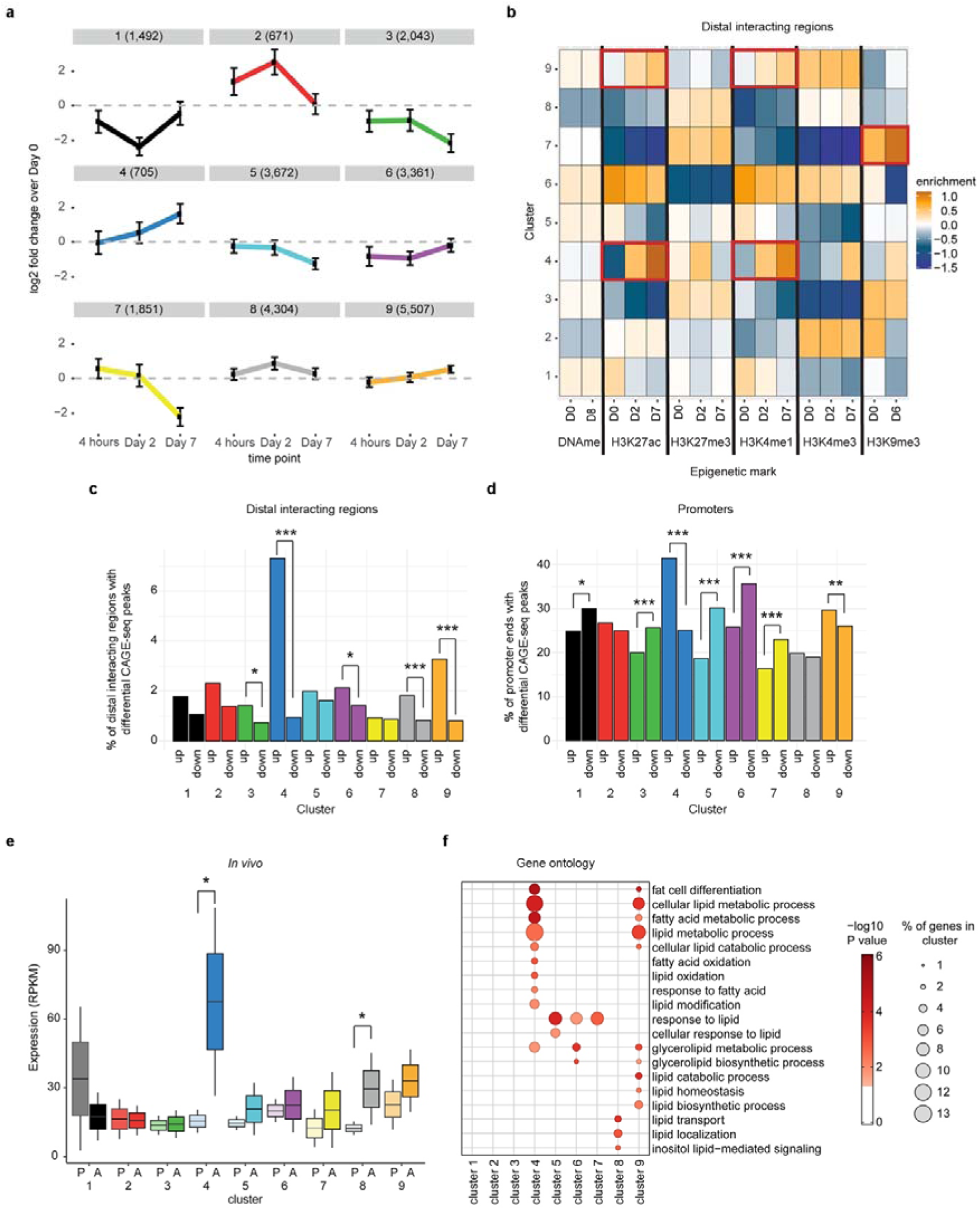
Cluster analysis of differential promoter-anchored interactions. a, The nine clusters of differential interactions (DInts) identified based on their dynamics during D0, 4 h, D2, and D7 stages of 3T3-L1 adipogenesis. Numbers in parentheses indicate the number of DInts per cluster. Interaction frequencies are shown as log₂ fold changes relative to D0 (horizontal dotted lines). Clusters 3, 4, 5, 7, and 9 show significant changes during the D2–D7 transition. Error bars represent s.d. b, Dynamics of epigenetic marks at distal interacting regions involved in DInts. Enrichment is relative to the average level of each modification across the nine clusters. c, Percentages of distal interacting regions per cluster containing differential CAGE-seq peaks. Clusters 3, 4, 6, 8, and 9 are enriched for CAGE-seq peaks upregulated at D7, consistent with enhancer activity. *P < 0.05, ***P < 0.001 by Fisher’s exact test. d, Percentages of promoters per cluster containing differential CAGE-seq peaks. Clusters 4 and 9 are enriched for peaks upregulated during adipocyte differentiation, while clusters 1, 3, 5, 6, and 7 are enriched for downregulated peaks. *P < 0.05, ***P < 0.001 by Fisher’s exact test. e, Genes from clusters 4 and 8 exhibit higher expression in adipocytes compared to preadipocytes in vivo, suggesting enrichment for adipogenesis regulators. Data are presented as box plots with mean (horizontal line) and 95% confidence intervals (error bars). *P < 0.05 by paired t-test. P, preadipocyte; A, adipocyte. f, Dot plot showing Gene Ontology (GO) terms related to adipocyte differentiation and lipid metabolism enriched in genes from DInts clusters. Dot colour and size represent significance and proportion of genes per cluster, respectively. See Supplementary Table 2e for a full list of significantly enriched GO terms.

To gain mechanistic insights into how these chromatin interaction dynamics relate to epigenetic regulation, we analysed histone mark profiles at both promoter and distal interacting regions of all nine DInt clusters. We used previously published ChIP-seq data for four histone modifications: H3K27ac and H3K4me1 (enhancer-associated), H3K27me3 (repressive), and H3K4me3 (active promoter-associated), collected at D0, D2, and D7^16^. In addition, we integrated DNA methylation profiles^23^ and H3K9me3 ChIP-seq data^24^ to evaluate broader chromatin state transitions (Fig. 2b; Extended Data Fig. 2a). The most striking changes occurred at distal regions of cluster 4, which gained H3K4me1 and H3K27ac and lost H3K27me3 between D2 and D7—consistent with enhancer activation. Cluster 9 showed a similar, but milder pattern. Distal regions in clusters 3 and 5 had modest loss of H3K4me1 and H3K27ac, while those in cluster 7 gained H3K9me3. In contrast, promoter-associated histone marks remained relatively stable across all clusters (Extended Data Fig. 2a). DNA methylation varied between clusters but was globally stable across the two time points (Fig. 2b; Extended Data Fig. 2a). We then examined dynamic transcriptional activity within DInt clusters by quantifying CAGE-seq peaks. Distal regions in cluster 4 showed the strongest enrichment for upregulated CAGE-seq peaks during adipogenesis, followed by clusters 3, 6, 8, and 9 (Fig. 2c). Promoters in clusters 4 and 9 were also enriched in upregulated peaks, whereas those in clusters 1, 3, 5, 6, and 7 were enriched for downregulated peaks (Fig. 2d).

To assess in vivo relevance, we analysed RNA-seq data from GFP^+^ pre-adipocytes and mature adipocytes isolated from Zfp423^GFP^ mice. Clusters 4 and 8 genes showed significant upregulation in adipocytes compared to pre-adipocytes (Fig. 2e). Analysis of published RNA-seq data from 3T3-L1 cells^16^ further confirmed that genes in clusters 4, 8, and 9 were significantly upregulated between D2 and D7, consistent with their chromatin rewiring patterns (Extended Data Fig. 2b; Supplementary Table 2c).

We next used i-cisTarget^25^ to identify transcription factor (TF) motif enrichment across promoter and distal regions of the nine clusters (Extended Data Fig. 2c; Supplementary Table 2d). Distal regions had markedly more TF binding motifs, many of which were exclusive to individual clusters. Notably, cluster 1 distal elements were enriched for CEBPA, CEBPB, CEBPD, and CEBPE motifs; cluster 3 for GLI1–3; cluster 4 for KLF6 and KLF13; cluster 5 for PLAGL1; and cluster 9 for PML, SIN3A, and ZBTB33. This cluster specificity suggests differential recruitment of TFs to enhancers guiding stage-specific gene programs. In contrast, promoter regions were enriched for broadly acting TFs shared across multiple clusters, including NRF1 (clusters 3, 4, 6, 8, 9), E2F4 (clusters 1, 3, 6, 8, 9), and ELF1 (clusters 5–7, 9).

Pathway enrichment analyses revealed functional distinctions among clusters. Clusters 4 and 9 were enriched in genes involved in fat cell differentiation (GO:0045444) and lipid metabolism (GO:0006629) (Fig. 2f; Supplementary Table 2e). Clusters 5, 6, and 7 were enriched for lipid response genes (GO:0033993), while cluster 8 showed enrichment in lipid transport (GO:0006869) and localization (GO:0010876). Clusters 1 and 3 were associated with gene expression regulation and developmental processes. Cluster 2 lacked significant GO term enrichment (Fig. 2f; Supplementary Table 2e).

In summary, our temporal clustering approach revealed distinct patterns of promoter-anchored chromatin rewiring throughout adipogenesis. Cluster 4 interactions, in particular, became markedly stronger between D2 and D7 and were associated with enhancer activation, transcriptional upregulation, and in vivo gene induction during late-stage adipogenesis. These results emphasize the importance of dynamic long-range chromatin interactions in orchestrating terminal differentiation programs and highlight enhancer–promoter communication as a central regulatory mechanism in the later stages of adipocyte maturation.

### siRNA knockdown of DInt-associated genes identifies novel regulators of late-stage adipogenesis

To investigate the functional relevance of genes involved in DInts, we conducted a genome-wide siRNA screen using the Mouse siGENOME Druggable Genome Library, targeting 2,905 genes (Supplementary Table 3a). The screen was performed in OP9-K mouse stromal cells^26^ (female; see Methods), a model that enables rapid adipocyte differentiation in vitro. Genes were considered hits if siRNA knockdown caused ≥30% change in lipid droplet formation (Extended Data Fig. 3a). This screen identified 981 genes whose knockdown reduced lipid accumulation, and 41 whose knockdown increased it (Fig. 3a). Known regulators of adipogenesis^1^, including Pdgfrb and Pparg, were among the hits (Supplementary Table 3a). Of the 2,905 genes, 1,216 were part of the nine DInt clusters. Among these, 432 knockdowns significantly altered lipid accumulation (410 reduced, 22 increased; Supplementary Table 3b). Notably, only cluster 4 was enriched for genes whose knockdown increased lipid accumulation (Fig. 3b), consistent with its late-stage enhancer activation.

**Fig. 3:**
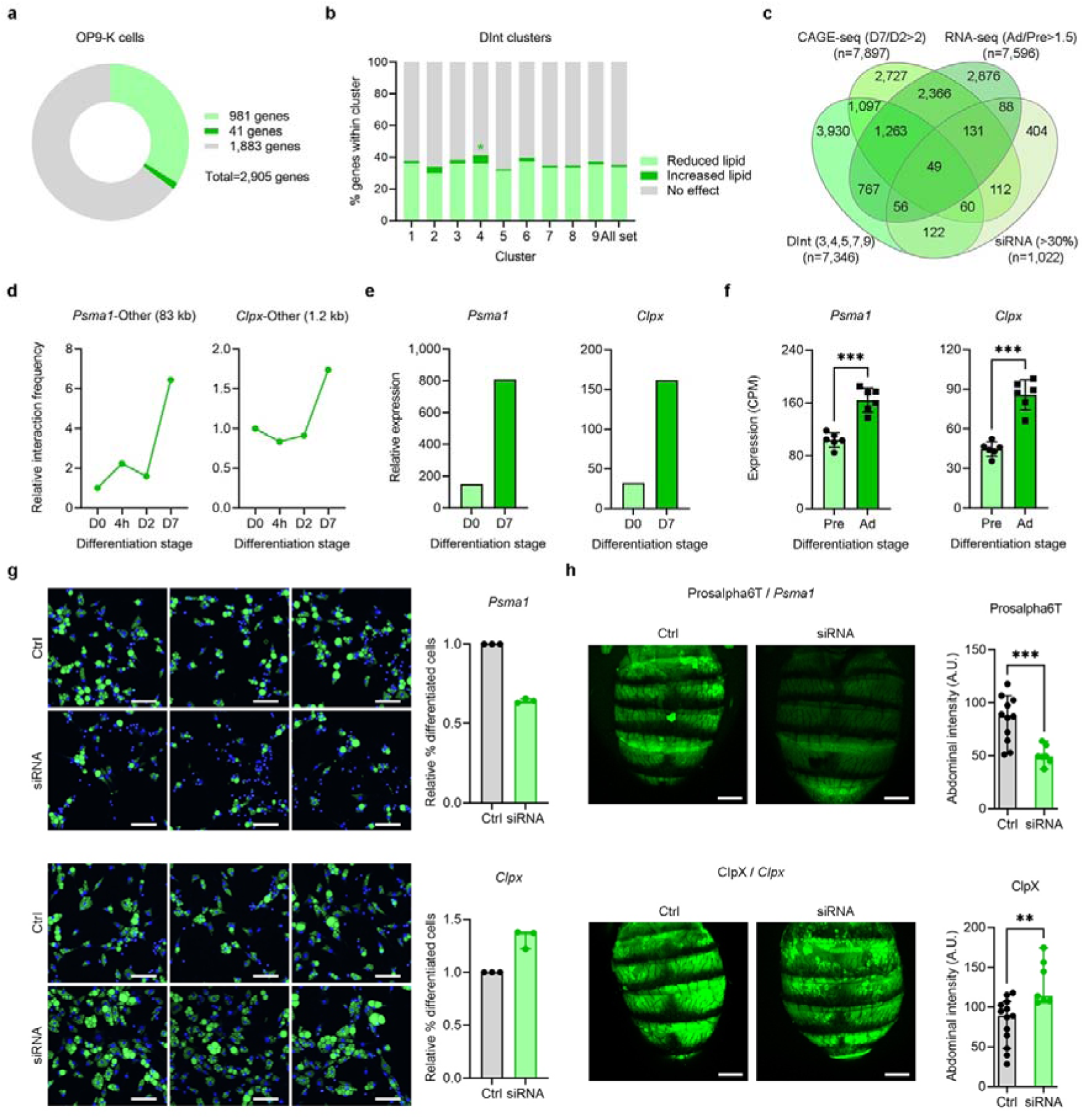
Validation of novel regulators of late-stage adipogenesis by siRNA knockdowns in vitro and in vivo. a, Overview of siRNA screen results in OP9-K cells: light green indicates >30% reduction in lipid accumulation, dark green indicates >30% increase, and grey indicates <30% change. b, Proportions of genes per DInts cluster that alter lipid accumulation upon siRNA knockdown in OP9-K cells. Cluster 4 is significantly enriched for genes that increase lipid accumulation compared to all 2,905 genes screened (*P < 0.05, two-sided χ² test). c, Venn diagram showing 49 candidate regulators of late adipogenesis fulfilling all four selection criteria (see main text). d, Dynamics of Psma1 and Clpx differential interactions during 3T3-L1 differentiation, quantified by GOTHiC. e, Relative expression of Psma1 and Clpx in 3T3-L1 preadipocytes (D0) and adipocytes (D7), measured by CAGE-seq. f, Relative expression of Psma1 and Clpx in primary mouse preadipocytes and adipocytes, measured by RNA-seq (CPM, counts per million). Error bars represent s.d.; ***P < 0.001. g, Left: representative images of OP9-K adipocytes stained with Bodipy (lipid droplets, green) and DAPI (nuclei, blue) following siRNA knockdown of Psma1 and Clpx (scale bars, 50 µm). Right: quantification of the percentage of differentiated cells relative to control (Ctrl), normalized to 1. Error bars represent s.d. h, Left: representative Bodipy-stained images of Drosophila abdomens following Prosalpha6T and ClpX siRNA knockdown (scale bars, 200 µm). Right: quantification of Bodipy intensity in the fat body. Error bars represent s.d.; **P < 0.01, ***P < 0.001 by unpaired t-test with Welch’s correction.

To identify novel regulators of late adipogenesis with high confidence, we applied an intersectional filter comprising four criteria: (1) genes with siRNA knockdown effects on lipid accumulation (n=1,022); (2) genes with promoters participating in DInts showing the largest changes between D2 and D7 (clusters 3, 4, 5, 7, and 9, representing 13,778 DInts involving 7,346 promoters); (3) genes with ≥2-fold expression change by CAGE-seq between 3T3-L1 pre-adipocytes and adipocytes (n=7,807); and (4) genes with ≥1.5-fold change in expression by RNA-seq between primary pre-adipocytes and adipocytes (n=7,596). This integrative approach yielded 49 genes involved in 122 DInts (Fig. 3c; Supplementary Table 3c). Manual curation (see Methods) identified 19 of these as previously unlinked to adipogenesis (Extended Data Fig. 3b), including genes encoding peptidases (e.g., Cela1, Lap3, Prss23, Clpx) and ubiquitin ligases (e.g., Znrf2, Rnf125, Rnf139, Hectd2, Fbxo17, Fbxl14, Trim21) (Supplementary Tables 3d–f).

We next evaluated conservation and function of these genes in Drosophila melanogaster, a well-established model for adipose biology^27^. Based on FlyBase homology scores (>12/15), 13 of the 49 and 5 of the 19 genes had strong fly homologs (Supplementary Tables 3c,f). Using the fat body-specific ppl-GAL4 driver^28^, we performed in vivo RNAi knockdowns during larval development. The fat body, the primary lipid storage and metabolic organ in flies, is functionally comparable to mammalian adipose tissue and liver. We selected three genes for in vivo validation: two novel candidates (Clpx, Fbxl14) and one known regulator (Psma1) as a positive control. In adult flies, fat body-specific knockdown of Psma1 (homolog Prosalpha6T) and Clpx led to lipid accumulation changes consistent with OP9-K results (Fig. 3d–h, Extended Data Fig. 3d–f), while Fbxl14 knockdown produced the opposite effect (Extended Data Fig. 3e–g), suggesting context-specific roles.

To assess clinical relevance, we examined whether genetic variation in the 19 novel genes is associated with human metabolic traits. We interrogated large-scale GWAS (n > 1.2 million individuals; see Methods) for five traits: body mass index (BMI), waist-hip ratio adjusted for BMI (WHRadjBMI), body fat percentage, triglyceride levels, and type 2 diabetes (T2D). Using variant-to-gene mapping^29^, incorporating eQTLs and activity-by-contact enhancer maps (see Methods), we linked six genes to proximal GWAS signals (Supplementary Table 4). FBXO17, FN3KRP, and ZFAND6 were associated with T2D (Fig. 4a), while FXYD5, LAP3, and SGPP1 were linked to circulating triglyceride levels (Fig. 4b). Notably, FN3KRP and ZFAND6 were the nearest genes to their lead SNPs, and the associated variants overlapped putative enhancers regulating their expression.

**Fig. 4:**
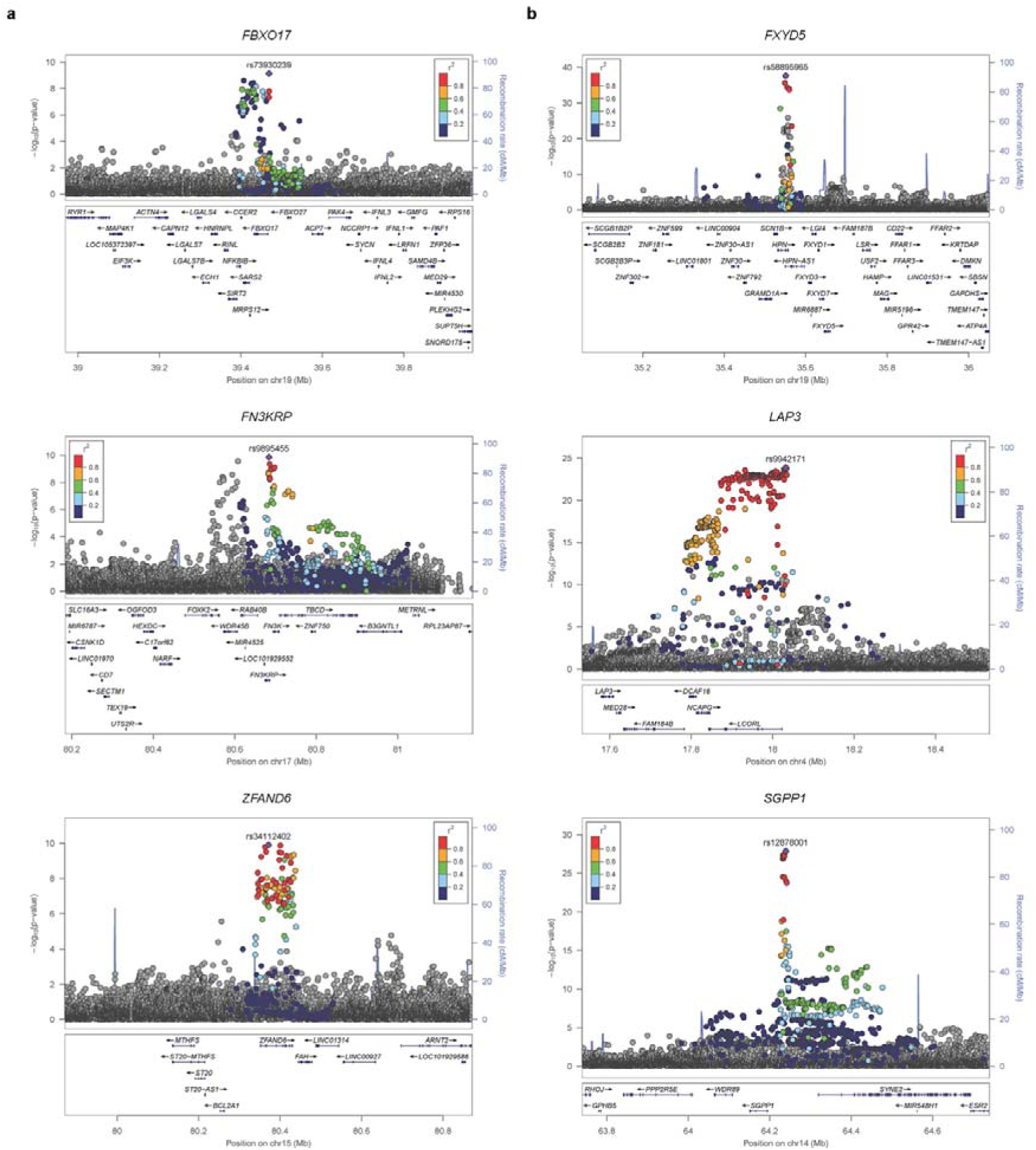
Novel regulators of late-stage adipogenesis associated with obesity and diabetes traits in human datasets. a, Regional association plots of GWAS signals near FBXO17 (rs73930239), FN3KRP (rs9895455), and ZFAND6 (rs34112402), identified as loci associated with increased risk of type 2 diabetes (T2D). b, Regional association plots of GWAS signals near FXYD5 (rs58895965), LAP3 (rs9942171), and SGPP1 (rs12878001), identified as loci associated with triglyceride levels. SNPs in linkage disequilibrium (LD; r² > 0.05) with each sentinel variant are color-coded according to the r² scale. Local recombination rates are shown as blue lines.

To explore functional links between these genes and adipose tissue biology, we analysed the Adipose Tissue Knowledge Portal^30^, which compiles transcriptomic and phenotypic data from over 6,000 individuals. Expression levels of FBXO17, FN3KRP, and ZFAND6 in subcutaneous adipose tissue negatively correlated with circulating insulin, WHR, and adipocyte volume, similar to PPARG (Extended Data Fig. 4). Conversely, expression of FXYD5, LAP3, and SGPP1 was positively correlated with BMI and adipocyte volume, mirroring patterns seen for LEP (Extended Data Fig. 4).

In summary, our integrated functional and genomic analyses identified 19 novel regulators of late-stage adipogenesis. These include genes encoding proteolytic enzymes and ubiquitin ligases, some of which are genetically linked to human metabolic traits. Validation across in vitro, in vivo, and large-scale human clinical datasets underscores their potential relevance to adipocyte biology and metabolic disease.

### Integrative mouse-human analysis links adipogenic chromatin rewiring to obesity and diabetes GWAS signals

To assess the relevance of DInt clusters to metabolic phenotypes, we examined two mouse mutant phenotype databases: MGI (13,551 genes with gain- and loss-of-function mutations) and IMPC (8,901 genes with loss-of-function mutations; 7,534 overlapping with MGI; see Methods and Supplementary Table 5). We focused on five phenotypes relevant to adipocyte biology: increased fat mass, decreased fat mass, abnormal fat morphology, altered circulating lipids, and insulin resistance/diabetes. In MGI, cluster 4 genes exhibited the highest odds ratios across all five phenotypes (Extended Data Fig. 5a; Supplementary Table 5b). Clusters 3, 5, and 9 also showed elevated odds for all five traits, while other clusters were associated with subsets of phenotypes. In IMPC, cluster 9 was enriched for increased fat mass; clusters 3 and 5 for decreased fat mass; and clusters 5 and 9 for abnormal fat morphology (Extended Data Fig. 5b; Supplementary Table 5c).

We next examined evolutionary conservation of distal interacting regions involved in adipogenesis-associated DInts. Using comparative genomics, we searched for human homologous regions with preserved synteny and trait-associated variation. The analysis had two steps: (1) identification of mouse DInts that met four criteria—conserved distal fragment, conserved gene, synteny between regions, and proximity to GWAS SNPs for obesity-related traits (Fig. 5a); (2) GWAS-to-gene (G2G) mapping^29^ to identify likely human causal genes. This integrative analysis yielded 596 SNP–trait associations implicating 249 candidate genes (Supplementary Table 6a). Of these, 31 genes were associated with two or more traits, and TCF7L2, a well-established T2D risk locus, was associated with all five traits (Fig. 5b; Supplementary Table 6b). Sixteen of the candidate genes also reduced lipid accumulation upon siRNA knockdown in OP9-K cells, providing orthogonal validation (Extended Data Fig. 6a).

**Fig. 5:**
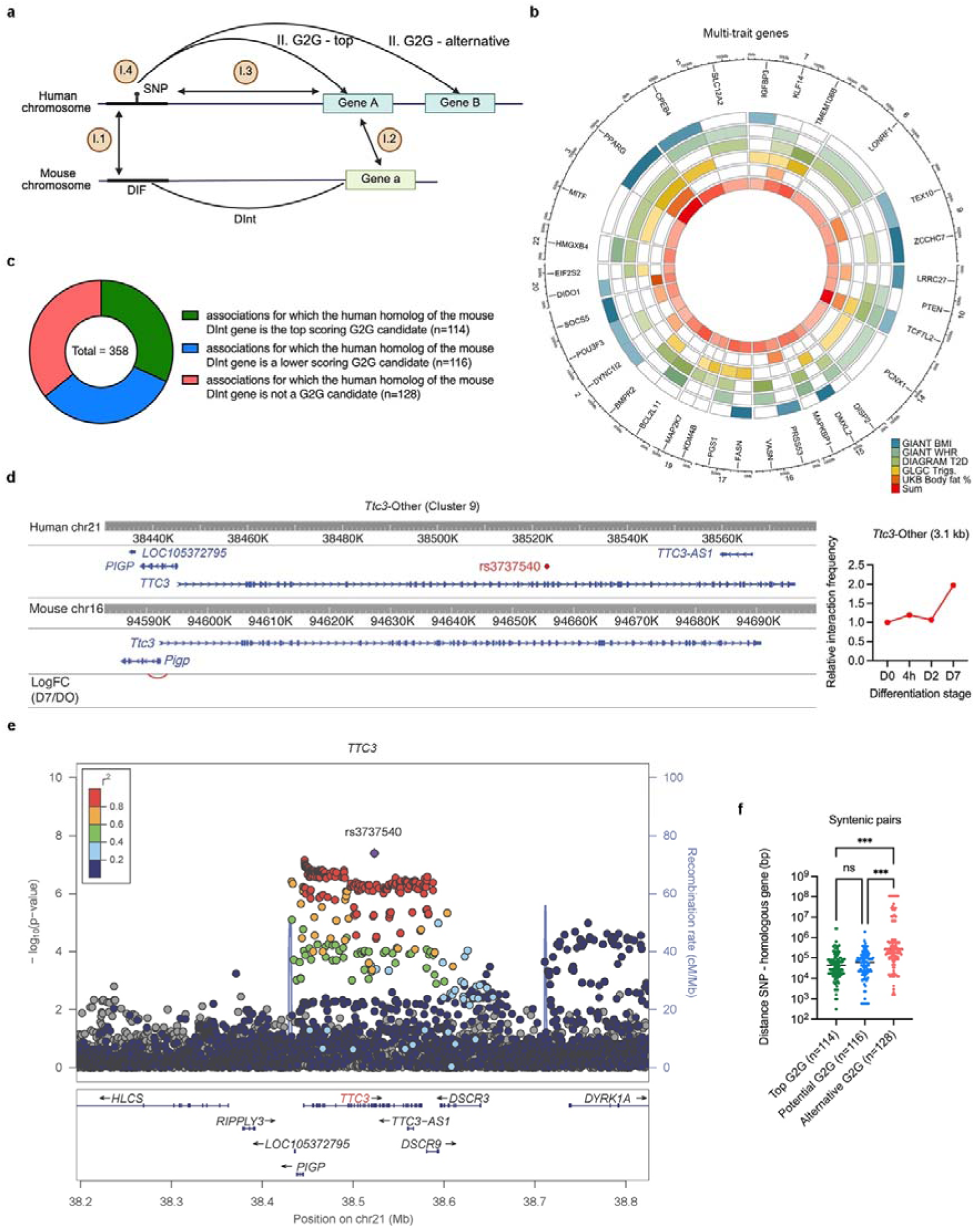
Validation of novel regulators of late adipogenesis using mouse–human synteny and human GWAS gene prioritization. a, Schematic overview of the two-stage analysis strategy. Stage I: mouse–human synteny analysis using four predefined criteria. Stage II: GWAS-to-genes (G2G) prioritization to identify the most likely causal gene (top-scoring) or lower-ranking candidate genes (alternative). b, Circular ideogram showing the 31 genes associated with at least two traits of interest. Trait categories are color-coded; gene positions on human chromosomes are shown on the outer ring. c, Donut chart summarizing the 358 SNP–trait associations that map to regions with conserved human–mouse synteny and overlap with differential interactions (DInts) from late adipogenesis clusters in 3T3-L1 cells. d, Left: synteny analysis between a region on human chromosome 21 (containing rs3737540, associated with type 2 diabetes in G2G analysis and implicating TTC3 as the candidate gene) and the homologous region on mouse chromosome 16 involved in a cluster 9 DInt. Right: dynamics of the cluster 9 DInt (Ttc3–Other) during 3T3-L1 differentiation, quantified by GOTHiC. e, Regional association plot of the GWAS signal near TTC3 (rs3737540), identified as a novel locus for T2D risk. SNPs in linkage disequilibrium (r² > 0.05) with the sentinel variant are color-coded according to r². Recombination rates are indicated by the blue line. f, Distance distribution between sentinel GWAS SNPs and G2G-prioritized genes. Comparisons are based on matches between the top G2G gene for each of the 358 traits and the human ortholog of mouse genes implicated in DInts. Data are shown as individual points with median and 95% confidence intervals. ns, not significant; ***P < 0.001 by Kruskal–Wallis test followed by Dunn’s multiple comparisons test.

Among the 596 associations, 358 involved DInts from clusters that show dynamic changes during late adipogenesis (clusters 3, 4, 5, 7, 9; Supplementary Table 6c). In 114 of these, the top G2G gene matched the human homolog of the mouse DInt gene (Fig. 5c; Supplementary Table 6d). These included 13 novel associations not yet listed in the GWAS Catalog—five for triglyceride levels (e.g., CDC148, IGFBP1, SYNCRIP) and eight for T2D (e.g., DISP2, EIF2S2, FAM53B, GNAO1, HMGXB4, TTC3).

Among these, TTC3 achieved the maximum G2G score (100th percentile) and was supported by all lines of evidence. Ttc3 knockdown reduced lipid accumulation in OP9-K cells (Extended Data Fig. 6a), and its promoter was involved in a cluster 9 DInt with a distal element ∼3.1 kb downstream (Fig. 5d). The homologous human region is in strong linkage disequilibrium (R² = 0.9) with the T2D-associated SNP rs3737540, which maps to an enhancer near TTC3 (Fig. 5d,e). In the Adipose Tissue Knowledge Portal^30^, TTC3 expression in subcutaneous fat negatively correlated with HOMA-IR, insulin, glucose, and fat cell volume (Extended Data Fig. 7).

Our analysis also refined existing GWAS assignments. For instance, the SNP rs1294410 (chr6) has been mapped to LY86 and BTF3P7 for WHRadjBMI^31^. Here, we implicate RREB1 as the likely causal gene (Supplementary Table 6d). This SNP lies within a mouse cluster 5 DInt with the Rreb1 promoter (Extended Data Fig. 6b), and Rreb1 expression is upregulated during adipogenesis in vitro and in vivo (Extended Data Fig. 6c). Although not tested in the OP9-K screen, Rreb1 has a conserved Drosophila homolog (peb; FlyBase score 12/15). Knockdown of peb in the fly fat body significantly reduced lipid content (Extended Data Fig. 6d). IMPC data revealed decreased fat content in males heterozygous for null alleles of Rreb1 and Ly86 (Extended Data Fig. 6e). Notably, these findings indicate that RREB1, rather than LY86 or BTF3P7, is the likely causal gene mediating the association between rs1294410 and WHRadjBMI.

In another 116 of the 358 SNP–trait associations (∼32%), the mouse DInt gene was not the top G2G hit, but ranked as a lower-confidence candidate (Fig. 5c; Supplementary Table 6e). Examples include AGRP for WHR (top gene: LRRC36), CKB for fat percentage (top gene: KLC1), and DOCK7 for triglycerides (top gene: ANGPTL3). In the remaining 128 associations, the DInt gene was not nominated by G2G (Supplementary Table 6f). These cases had significantly greater distances between the SNP and gene, with 27 located >1 Mb away—outside the G2G mapping window (Fig. 5f). For example, rs72926932 (chr18) is associated with T2D and mapped to TCF4^32^, similar to the GWAS Catalog. However, the homologous mouse region forms a cluster 9 DInt with the Rnf125 promoter, separated by 9.3 kb. In humans, the syntenic regions are >23 Mb apart (Extended Data Fig. 6f). Notably, Rnf125 knockdown in OP9-K cells reduced lipid accumulation (Extended Data Fig. 6a), and Rnf125 is one of the 19 novel regulators of late adipogenesis identified in this study. Thus, despite the distance, RNF125 may contribute to the observed T2D association, alone or alongside TCF4.

In summary, by integrating chromatin interaction dynamics, interspecies conservation, variant-to-gene mapping, and functional screens, we provide evidence that several DInt-associated genes modulate adipogenesis and are linked to human metabolic traits. This approach not only refines causal gene assignments at known loci but also uncovers novel candidate genes underlying susceptibility to obesity and type 2 diabetes.

## Discussion

The mechanisms that uncouple adiposity from cardiometabolic comorbidities are complex, involving genetic and environmental influences^33^. Both a deficit and an excess of fat cells—especially hypertrophic adipocytes—are associated with impaired metabolic health^7^. Recent studies have highlighted genetic and epigenetic determinants of limited adipose storage capacity^8,34^ and regional fat distribution^35,36,37,38^ that modulate insulin resistance risk, supporting the idea that subtle forms of lipodystrophy contribute to disease^39^. These findings underscore the importance of molecular mechanisms regulating late-stage adipogenesis in metabolic health, which remain incompletely understood.

Long-range chromatin interactions between promoters and distal regulatory elements play a critical role in transcriptional regulation during differentiation^40,41,42,43^, including early and intermediate stages of adipogenesis^14,44^. Our analysis of promoter-anchored chromatin loops that change between day 2 and day 7 of differentiation revealed dynamic remodelling of distal elements, many of which correspond to enhancer-like regions marked by increased H3K27ac and H3K4me1, active enhancer RNA transcription, and enrichment in genes upregulated during adipogenesis. Conversely, some distal elements gained repressive marks such as H3K9me3, potentially contributing to decreased chromatin looping and gene repression^45,46^. Transcription factor motif enrichment analyses showed stage-specific regulatory signatures, with early adipogenesis-associated clusters enriched for CEBPB and CEBPD motifs^4^, and late-stage clusters showing enrichment for KLF6^47^ and KLF13^48^, highlighting coordinated promoter and enhancer remodelling as a key driver of adipogenic transcriptional programs. Additionally, our CAGE-seq data revealed widespread promoter switching during differentiation, affecting over 600 genes, a mechanism known to diversify gene regulation in developmental transitions and cell type–specific functions^49,50,51,52,53^. Together, these findings underscore the intricate interplay between chromatin architecture and transcriptional control in adipocyte differentiation.

Building on the regulatory framework of chromatin dynamics, we identified 19 regulators of late-stage adipogenesis by integrating multi-omics data, siRNA screening of druggable genes, in vivo Drosophila validation, and cross-species GWAS and mutant phenotype analyses. Many of these 19 novel regulators are ubiquitin protein ligases, particularly genes encoding SCF (Skp1–Cullin1–F-box) complex components such as Znrf2, Rnf125, Rnf139, Hectd2, Fbxo17, Fbxl14, Trim21, and Fbxw5. Other E3 ligase-related genes included Traf2—which regulates NF-ĸB and JNK signalling^54^—and Zfand6, involved in mitophagy via TRAF2–cIAP complexes^55^. Except for Znrf2, knockdown of all ubiquitin ligase genes significantly reduced lipid accumulation in OP9-K cells, implicating the ubiquitin–proteasome system in terminal adipogenesis. In addition to ligases, several proteases emerged as novel regulators, including CLPX (a mitochondrial protease subunit involved in the unfolded protein response), PRSS23 (a secreted serine protease potentially regulating IGF binding proteins), CELA1 (which hydrolyses extracellular proteins such as elastin), and LAP3 (a cytosolic aminopeptidase implicated in redox and glutathione metabolism). These findings expand the landscape of proteostasis mechanisms, highlighting not only protein degradation but also regulated proteolysis as essential to terminal adipogenesis.

Population-level genetic and clinical datasets, including the Adipose Tissue Knowledge Portal^30^, confirmed the involvement of six of the 19 novel adipogenesis regulators—FBXO17, FN3KRP, FXYD5, LAP3, SGPP1, and ZFAND6—in human adipose-related traits. Further refinement through cross-species synteny analysis of conserved promoter–enhancer interactions pinpointed additional candidates such as TTC3 and RREB1. RREB1 has been associated with visceral fat accumulation in humans^56^, and recent studies demonstrate that its loss impairs adipogenesis while enhancing insulin sensitivity in murine and human cells^57^. TTC3, notable for its role in Akt degradation^58^, also functions as an E3 ubiquitin ligase, reinforcing the central involvement of ubiquitin-mediated pathways in adipocyte biology. Importantly, some conserved regulatory elements exhibit species-specific target gene interactions, exemplified by a region engaging mouse Rnf125 and human TCF4, illustrating regulatory repurposing across evolution. This integrative framework—combining functional genomics, evolutionary conservation, and human genetics—provides critical mechanistic insights into adipogenesis and obesity risk, addressing a major gap in the GWAS Catalog by attributing biological function to previously ambiguous genetic associations.

Our study has several limitations. Genetic influences on human adiposity vary by fat depot, age, and sex^37,59,60,61,62,63,64,65^, factors not modelled in the cell lines used here. Additionally, some dynamic promoter–enhancer interactions identified in mouse cells may not be conserved in humans due to chromosomal rearrangements. Future application of promoter capture Hi-C to human adipocyte differentiation models will be essential to validate and extend these findings.

In summary, our study provides compelling evidence that dynamic promoter–enhancer chromatin interactions orchestrate transcriptional programs underlying late-stage adipogenesis. We identified novel, druggable regulators—particularly E3 ubiquitin ligases and proteases—involved in proteostasis, whose roles were supported by siRNA screening, cross-species analyses, and genetic association data. These findings underscore the central role of distal regulatory elements in adipocyte maturation and offer a comprehensive framework integrating chromatin architecture, multi-omics, functional screens, and genetics to uncover novel regulators with relevance to human metabolic disease.

## Methods

### 3T3-L1 adipocyte differentiation

Mouse 3T3-L1 cells were grown at 37°C and 5% CO2 in Dulbecco’s Modified Eagle’s Medium (DMEM high glucose, Sigma-Aldrich #D6546) supplemented with 10% calf serum (Gibco #16010159), 2% glutamine (Sigma-Aldrich #G7513) and 1% Pen-Strep (Sigma-Aldrich #P0781). Cells were induced to differentiate two days post confluency (defined as day zero – D0) by addition of differentiation media: DMEM high glucose supplemented with 10% fetal bovine serum (Sigma-Aldrich #F7524), 1% Pen-Strep, 1 μg/ml insulin (Sigma-Aldrich #I9278), 390 ng/ml dexamethasone (Sigma-Aldrich #D4902), 1,115 μg/ml 3-Isobutyl-1-methylxanthine (Sigma-Aldrich #I5879) and 2 μM rosiglitazone (Sigma-Aldrich #R2408). Two days after the induction of differentiation, fresh differentiation media supplemented with 1 μg/ml insulin was added. From day four and onward, cells were maintained in differentiation media.

To quantify efficiency of 3T3-L1 differentiation into adipocytes, staining with Oil red O was performed, as previously described^66^. Briefly, cells were fixed in 10% formaldehyde in phosphate buffer (PBS) for one hour, washed with 60% isopropanol, and stained with Oil Red O solution (Sigma-Aldrich #102419) for 10 minutes followed by four washing steps with water, counterstaining of nuclei with hematoxylin (Sigma-Aldrich #105175) and imaging with an optical microscope. Counting the percentage of cells containing Oil Red O positive lipid droplet was performed by ImageJ software (National Institutes of Health, Bethesda MD, USA). Over 90% of cells were Oil Red O positive after seven days of differentiation, using the above protocol.

We verified 3T3-L1 differentiation efficiency by measuring mRNA levels of known markers of adipogenesis by qRT-PCR. Total RNA was extracted from D0 and D7 3T3-L1 cells using an RNeasy Plus Mini Kit (Qiagen #74134). RNA concentration was measured by NanoDrop (Thermo Fisher Scientific) and quality was assessed in agarose gels. Reverse transcription was performed using the RevertAid RT Reverse Transcription Kit (Thermo Fisher Scientific – K1622). qRT-PCR was performed with the SYBR Green JumpStart Taq Ready Mix (Sigma – S4438) and custom-made primers (Supplementary Table 7) using an ABI Prism 7900 system (Applied Biosystems). For gene expression normalization, we used two housekeeping genes (Ppia and Rplp0). Levels of expression were calculated using the 2^-ΔΔCt^ method.

### CAGE-seq analysis of D0 and D7 3T3-L1 cells

CAGE-seq analysis was performed using the nAnT-iCAGE method, as described^18^. Briefly, total RNA was extracted from four D0 and four D7 biological replicates of 3T3-L1 cells, using an RNeasy Plus Mini Kit (Qiagen #74134). RNA concentration and quality were measured by NanoDrop (Thermo Fisher Scientific) and an Agilent RNA 6000 Nano Kit (Agilent #5067-1511), respectively. All samples had RIN>9. Library preparation was performed at the Genome Network Analysis Support Facility, RIKEN CLST, with 50-base single read sequencing performed on an Illumina HiSeq2500 instrument. Reads were mapped using TopHat, and RECLU^67^ was used to call differentially expressed top peaks and bottom peaks corresponding to reproducible TSSs. Gene level expression was calculated from the RECLU output using top peaks. First, D0 peak level expression was obtained using the formula (2^logConc*2)/(1+2^logFC), then D7 expression values were calculated by D0*2^logFC. The expression values of top peaks annotated to the same gene were summed up for gene level expression. Genes with more than two differential top peaks, where the direction of change was divergent across the differential peaks, were identified as genes involved in promoter switch. Functional analysis was performed using DAVID (Database for Annotation, Visualization and Integrated Discovery; v6.8 https://david.ncifcrf.gov). Enriched gene ontology (GO) terms with FDR1<15% were considered significant. These terms were then clustered semantically using REViGO (Reduce and Visualize GO)^68^, which removes redundancy, and ordered according to the log_10_ p values.

### RNA-seq analysis of primary pre-adipocytes and adipocytes

Pre-adipocytes and adipocytes were isolated from epididymal fat pads of young (three months-old) hemizygous Zfp423^GFP^ male mice^19^. RNA was extracted using the RNeasy Plus Micro Kits (Qiagen #74034) and used to prepare RNA libraries utilizing the TruSeq Stranded mRNA Kit (Illumina #20020595) and sequenced as single-end 50 bp reads using an Illumina HiSeq 4000 platform.

RNA-seq reads were mapped to GRCm39 version of the mouse reference genome sequence using STAR (v2.5.1b)^69^. Reads were considered mapped if the similarity was greater than 95% over at least 90% of the read length, as previously described^70^. FeatureCounts (v1.5)^71^ was applied for the generation of count tables based on the mapping files. Raw counts were subjected to differential gene expression analysis via DESeq2^72^ and normalized to CPM (counts per million).

### Promoter capture Hi-C (PCHi-C) analysis

PCHi-C was performed in triplicate for D0 and D7 3T3-L1 cells as previously described^12^. Briefly, 30-40 million cells/sample were fixed with 2% formaldehyde (Agar Scientific #AGR1026) for 10 min at room temperature, prior to harvesting from the tissue culture flasks. Digestion of cross-linked chromatin with HindIII (NEB #R3104), labelling with biotin-14-dATP (Invitrogen #19524016) and DNA Polymerase I (Large Klenow Fragment; NEB #M0210) and ligation (with T4 DNA ligase; Invitrogen #15224025) were all performed on nuclei kept intact. Chromatin cross-linking was then reversed by incubation with proteinase K (Roche #03115879001) at 65°C overnight and the DNA was purified by phenol/chloroform extraction. Aliquots of the Hi-C libraries were used to verify efficiency of chromatin digestion and digestion using standard PCR and qPCR, with primers listed (Supplementary Table 7). Then, biotin from un-ligated DNA ends was removed using T4 DNA polymerase (NEB #M0203) and the DNA was sonicated (Covaris E220) to an average size of around 400 base pairs. Sonicated DNA was end-repaired using DNA Polymerase I (Large Klenow Fragment; NEB #M0210), T4 DNA polymerase (NEB #M0203) and T4 DNA polynucleotide kinase (NEB #M0201), then dATP was added to the 31 ends of the DNA using Klenow exo-(NEB #M0212). The DNA was then subjected to double-sided SPRI bead size selection (AMPure XP beads; Beckman Coulter #A63881) and biotin-marked ligation products were isolated using MyOne Streptavidin C1 Dynabeads (Invitrogen #65002). After adapter ligation (Illumina PE PCR 1.0 and PE PCR 2.0 primers), the bead-bound Hi-C DNA was amplified with seven PCR amplification cycles. Promoter capture Hi-C was performed using a custom-made RNA capture bait system (Agilent Technologies) consisting of 39,021 individual biotinylated RNAs targeting the ends of 22,225 promoter-containing mouse HindIII restriction fragments, as described^12^. After a post-capture PCR (four amplification cycles using Illumina’s PE PCR 1.0 and PE PCR 2.0 primers), the PCHi-C libraries were purified with AMPure XP beads (Beckman Coulter #A63881) and paired-end sequenced (HiSeq 1000, Illumina) at the Babraham Institute Sequencing Facility.

Quality control of raw fastq files was performed using FastQC (https://www.bioinformatics.babraham.ac.uk/projects/fastqc/) and trimming was done when required using TrimGalore (https://github.com/FelixKrueger/TrimGalore). Reads (220 – 282 million paired-reads/sample) were mapped to the mm9 genome and filtered using the HiCUP pipeline (v0.5.8), which removes experimental artefacts, such as circularised reads and re-ligations, and duplicated reads^73^. After de-duplication, the number of valid unique di-tags varied between 90 – 154 million per sample (i.e., between 92.8% and 99.5% of the valid pairs). GOTHiC^20^ was used to identify significant differential interactions (DInts) by comparing our own D0 and D7 data, as well as that obtained previously at D0, 4h and D2^14^. DInts were taken forward when they overlapped in at least two replicates and log_2_ fold changes were calculated as the average between the replicates in which the interaction was classified as DInts. Interactions overlapping with two regions of known artefacts (chr10:106613366-107858706 and chr10:116174799-118176364) were removed.

### q3C assays

3C (chromatin conformation capture) assays were performed in independent biological replicates of D0 and D7 3T3-L1 cells. Briefly, 10-15 million cells per sample (6-9 D0 samples and 7 D7 samples) were fixed with formaldehyde, digested with HindIII and ligated with T4 DNA ligase as described above, without the incorporation of biotin-14-dATP. After reversal of chromatin crosslinking, the purified DNA was used in qPCR reactions using 100 ng DNA as template/reaction, the SYBR Green JumpStart Taq Ready Mix (Sigma – S4438) and custom-made primers (Supplementary Table 7) using an ABI Prism 7900 system (Applied Biosystems). Interaction frequencies were calculated using the 2^-ΔΔCt^ method and normalized against the interaction between two consecutive HindIII fragments at the Rplp0 locus, used as internal control for efficiency of HindIII chromatin digestion and DNA ligation.

### Clustering of DInts

Interactions that were significant in at least one of the three comparisons (4h vs D0, D2 vs D0, D7 vs D0) were used for clustering. Log_2_ fold changes of the interactions were collected from all three comparisons and grouped using k-means clustering^74^. The number of clusters was determined using the elbow method by calculating the within group sum of squares for k=2 to k=15. The optimal number of clusters was 9. The profile of the log_2_ fold changes of differential interactions was visualized using ggplot2 showing the mean log_2_ fold changes and error bars indicating the standard deviation within the cluster.

### ChIP-seq and WGBS data processing and enrichment calculation

ChIP-seq data for H3K27ac, H3K27me3, H3K4me1 and H3K4me3 at D0, D2 and D7 was obtained from Mikkelsen et al.^16^. DNA methylation data was used from Park et al.^23^ and H3K9me3 from Matsumura et al.^24^. For H3K27ac, H3K27me3, H3K4me1 and H3K4me3 we made use of the peaks identified by Mikkelsen et al. The H3K9me3 ChIP-seq data quality control was performed using FastQC, and reads were trimmed using TrimGalore (https://github.com/FelixKrueger/TrimGalore). Mapping to the mm9 genome was achieved using BWA-MEM (<u>arXiv:1303.3997</u>) and broad peaks were called using MACS2^75^. We overlapped histone modification peaks with promoter or distal fragments belonging to the nine clusters or the differential interactions between D0 and D7 and labelled each fragment whether it carried the mark or not. Enrichment of fragments with the mark was calculated across clusters/categories and was normalized for the total number of peaks found for that modification at that time point. For DNA methylation, average methylation level was calculated for each HindIII fragment in the genome and enrichment in a cluster/category was compared to the average across them.

### Expression microarray analysis in 3T3-L1 and OP9-K cells

OP9-K is an embryonic stromal cell-line of unspecified sex (www.cellosaurus.org). To establish the sex of origin of these cells, we compared gene expression levels in OP9-K cells^26^ with those of male white adipose tissue (C3HeB/FeJ), as well as of male and female liver samples, retrieved using the accession numbers GSE197101 and GSE176226, respectively. Microarray data normalization was conducted using affy, oligo, and limma packages in R. Then, data were annotated using GEO reference table (https://www.ncbi.nlm.nih.gov/geo/query/acc.cgi?acc=GPL6246) to identify the chromosomal location of each gene. Normalized expression values were averaged for replicates and expression levels in the OP9-K cells were compared to the expression in male and female tissues across all chromosomes and separately on the Y chromosome. These analyses revealed that OP9-K cells originated from a female embryo.

3T3-L1 was reported as a spontaneously immortalized cell-line isolated from a male embryo (www.cellosaurus.org). However, recent karyotyping analyses revealed the presence of two X chromosomes^76^. To establish the sex of 3T3-L1 cells, microarray expression data retrieved using the accession number GSE20752 was normalized with other male and female data. Male white adipose tissue and female mammary gland expression data were taken from GSE10246. All the raw expression data were normalized using affy and limma packages in R. Normalized expression values were averaged across the replicates. Then, the probes were annotated using the GPL1261 annotation table from GEO (https://ftp.ncbi.nlm.nih.gov/geo/platforms/GPL1nnn/GPL1261/annot/) to identify and compare the expression level of genes located on each chromosome and specifically on chromosome Y between the 3T3-L1 cell lines and the known gender tissues. Our analyses are consistent with recent findings suggesting origin in a female embryo.

To compare mRNA expression levels at D0, D2 and D7 in 3T3-Le cells, data generated using GeneChip arrays (Affymetrix) were retrieved from Table S2 of Mikkelsen et al.^16^. Data was sorted per DInt cluster, with a coverage between varying between 59.2% of genes (cluster 7) and 71.8% (cluster 4). Data was analysed per individual DInt cluster using one-way ANOVA tests followed by Tukey’s multiple comparisons tests.

### TF enrichment analysis

Up- and down-regulated CAGE-seq peaks were overlapped with the promoter and distal fragments of differential interactions across the nine clusters. The positions of CAGE-seq peaks (>7bp) were used in i-cisTarget^25^ to test for enrichment of known motifs and TF binding based on publicly available ChIP-seq data. TFs with a normalized enrichment score (NES) above 5 were considered significant.

### Pathway analysis

Genes involved in differential interactions across the nine clusters were tested for enrichment of GO biological processes using g:Profiler^77^. The lists of significantly enriched (p-adj <0.05) GO terms were summarized and visualized with REVIGO^68^.

### siRNA screen for regulators of adipocyte differentiation in the OP9-K cells

Mouse OP9-K cells were grown at 37°C and 5% CO2 in OP9 propagation media: MEM-α (Life Technologies #12571-063), supplemented with 20% FBS (Sigma-Aldrich #F4135), 2mM L-glutamine (Sigma-Aldrich #G7513) and 1% Pen-Strep (Sigma-Aldrich #P0781). Cells were passaged at 80% confluence and fed every 2-3 days. The siRNA screen was performed using SMARTpool siRNAs (i.e. a pool of four siRNAs per gene) from the Mouse Druggable Subset consisting of 2,905 genes distributed on 11 384-well plates (Dharmacon #G-014675-E2, Lot 14008). The 0.1nmol lyophilised siRNA stock plates were re-suspended to 10µM, and then further diluted to give working stock plates of 3µM siRNA, which was used for transfection.

The siRNA screen was performed in 384-well round bottom plates coated with poly-D-lysine (Sigma-Aldrich #P7886). Briefly, 2.6µl of 3µM working siRNA was pipetted into each well, followed by 3µl of the transfection mix (obtained by mixing 339.6µl Lipofectamine RNAiMAX transfection reagent [Invitrogen #56532] + 4646µl OptiMEM [ThermoFisher Scientific #31985062]) and, after 5 minutes, 30µl cell suspension (diluted at 133 cell/µl). To ensure even distribution of cells into the wells, plates were incubated undisturbed at room temperature for 15 minutes before transferring carefully to the incubator at 37°C and 10% CO2 (shown to give better differentiation). Non-targeting ON-TARGETplus pool (Dharmacon #D-001810-10-20), mouse Pparg ON-TARGETplus SMARTpool and mouse L3mbtl3 ON-TARGETplus SMARTpool were used as controls on each 384-well plate. All pipetting was performed using and FXP robot and Biomek AP96 P200 pipette tips (Beckman Coulter #717252).

Differentiation was initiated 24 hours post-transfection using freshly prepared insulin-oleate media: MEM-α supplemented with 0.2% FBS, 175 nM insulin, 900 µM oleate:albumin (Sigma-Aldrich #O3008) and 1% Pen-Strep and cells were incubated for 48 hours undisturbed to allow adipocyte differentiation. Following differentiation, the cells were fixed for 20 minutes by adding 6µl 24% formaldehyde directly to medium in each well. After washing with PBS, cells were stained for 30 minutes with 42µl BODIPY 493/503 for lipids (1mg/ml in ethanol; ThermoFisher Scientific #D3922) and 42µl Hoechst 33342 for nuclei (10mg/ml; Invitrogen #H3570) and washed again in PBS.

The plates were scanned on the ImageXpress Confocal Micro (Molecular Devices) using the 10x objective and the confocal pinhole setting. Cells were imaged in two channels: the Hoechst channel to capture nuclei and the FITC channel to capture the lipid droplets. To correct for unevenness across the well, three images were taken at different z planes and a maximum projection image was compiled and used for all future analyses. Images were analysed using the Multiwavelength Cell Scoring Application Module within the MetaXpress software (Molecular Devices). Nuclei were defined in the Hoechst channel by size and intensity with user-defined thresholds. The cytoplasm was defined using the lipid stain and was thresholded by size and intensity. Cells were then classified as differentiated if the lipid-stained cytoplasm exceeded a defined minimum stained area of 100µm^2^ (Extended Data Fig. 3a). Intra-plate normalisation was carried out for each of the three triplicates, then the median of the three values for each well was calculated. This value, termed median relative differentiation (i.e. values normalised to non-targeting), was used to determine hits. Cuts-off of 0.7 (i.e. a 30% decrease in differentiation) and 1.3 (i.e. a 30% increase in differentiation) were used to identify genes that decrease and increase adipocyte differentiation, respectively.

### siRNA knockdowns in Drosophila fat body

All fly strains were maintained in Darwin Chambers (IN084-AA-LT-DA-MP) at a temperature of 25°C. and 70% humidity with 12h:12h light-dark cycles and reared on Nutri-Fly Bloomington Formulation food medium (Genesee Scientific #66-112). The fly lines used in this study were obtained from Bloomington Drosophila Stock Centre (BDSC): Ppl-Gal4 (BDSC #58768), UAS-Peb-RNAi (BDSC #33943) UAS-Ppa-RNAi (BDSC #31357), UAS-Prosalpha6T-RNAi (BDSC #55243) and UAS-Clpx-RNAi (BDSC #57577). RNAi-mediated knockdown specifically in the fat body was achieved by driving UAS-RNAi expression using the fat body specific driver line, Ppl-GAL4. Female flies from F1 generation bearing both Gal4 and UAS-RNAi constructs were used for fat body staining. F1 female progenies obtained from the cross between the Ppl-Gal4 and the UAS-RNAi lines with isogenic w1118 (BDSC #6326) wild-type flies were used as genotypic controls.

To verify the efficiency of knockdowns, total RNA was extracted from whole bodies of flies using TRI-reagent (Sigma #93289) and 2µg of total RNA was reverse transcribed using High-capacity cDNA synthesis kit (Applied Biosystems #4374967). Real-time qPCR was performed using PowerUp SYBR Green Master Mix (ThermoFisher #A25741) with listed primers (Supplementary Table 7). Each qRT-PCR reaction was performed in duplicate. Rpl32 was used as an endogenous control. The cut-off for Ct values was <35 for testing genes and <25 for Rpl32. Relative expression analysis was done using the 2^-ΔΔCt^ method. Six to eight independent biological replicates (containing 5 whole bodies each) per genotype were tested for each experiment.

Subcuticular fat body staining of undissected fly abdomens was adapted from Li et al.^78^. Briefly, 2-5 days-old female flies were anesthetized to remove the legs and wings. The fly bodies were then fixed in 4% paraformaldehyde for 20 minutes, followed by washing in phosphate-buffer saline (PBS). The flies were then submerged thrice in liquid Nitrogen for few seconds, each time followed by thawing on ice for 1 minute. A solution of 1µg/µL of BODIPY – 493/503 (1:500 dilution, ThermoFisher, #D3922) in PBS was added to the samples and incubated under dark for 1 hour, followed by three washes with PBS. The flies were then mounted onto a glass slide by gluing the thorax on the ventral side. The samples were covered with Vectashield mounting medium (Vector Laboratories #H-1000-10) and imaged by confocal microscopy (Nikon A1R) under the FITC channel. Same confocal setting was used across all samples. Maximum intensity projection images were analysed and quantified in ImageJ.

### Manual curation of 49-gene list to identify novel regulators of late-stage adipogenesis

This analysis was performed in two steps. First, three authors carried out an independent search on GWAS (https://www.ebi.ac.uk/gwas/home), OMIM (https://www.omim.org), IMPC (https://www.mousephenotype.org) and MGI (https://www.informatics.jax.org/) for evidence linking the gene of interest with obesity, lipodystrophy or other phenotypes associated with altered fat mass or adipocyte differentiation. This was supplemented with a search on PubMed (https://pubmed.ncbi.nlm.nih.gov) using a range of keywords (adipogenesis, adipocyte differentiation, lipodystrophy, obesity, adiposity, fat mass, body mass index) plus the gene symbol. The information was then verified and summarized by two other authors, leading to the identification of 19 genes without any prior evidence for a role in adipogenesis or related processes.

### Analyses of MGI and IMPC obesity and diabetes phenotypes

We retrieved the phenotyping data for 13,551 genes from MGI, using the batch query function (data accessed in April 2024). The phenotyping terms were grouped semantically in five categories: increased fat amount (n=31 mammalian phenotype – MP – terms), decreased fat amount (n=31 MP terms), abnormal fat morphology (n=33 MP terms), altered circulating lipid levels (n=25 MP terms) and insulin resistance/diabetes (n=12 MP terms) (Supplementary Table 5a). In the case of IMPC (which contains phenotypic data for 8901 knockout genes – data release version 21.1; accessed in July 2024), we downloaded the lists of genes related to five phenotypes of interest: increased total body fat amount (367 genes with significant changes), decreased total body fat amount (293 genes), abnormal adipose tissue morphology (653 genes), abnormal circulating triglyceride levels (229 genes) and abnormal circulating insulin levels (68 genes) (Supplementary Table 5d). For both datasets, the enrichment of genes with significant phenotypes within the nine clusters of DInts was calculated using Fisher’s exact tests and represented as odds ratios ± 95% confidence intervals. To verify the specificity of our findings, we also performed similar analyses for genes that do not belong to any of the nine DInts clusters and for two unrelated phenotypes (abnormal anxiety-related response – 143 genes, and abnormal heart morphology – 543 genes) in the case of MGI and IMPC data, respectively. None of these control searches associated any significant enrichment in the nine clusters of DInt genes (Extended Data Fig. 5; Supplementary Tables 5b,c).

### Mouse-human synteny analyses

We have assessed synteny based on synteny blocks identified using the synteny portal^79^ and defined the exact homologous positions from the UCSC hg19-mm9 syntenic Net files. Differential interactions, where both fragments fall into the same synteny block, and obesity-related intergenic, intronic and TF binding site SNPs falling into synteny blocks were identified. Those SNPs that overlapped with the distal fragment of a differential interaction were mapped to the gene involved in that interaction.

### Human GWAS gene prioritisation

The identified DInts and DInt-linked genes were mapped to syntenic regions in the human genome as described above and integrated with human genome-wide association study (GWAS) data on five relevant metabolic traits: body mass index (BMI) and waist-hip ratio (WHR) adjusted for BMI in up to 806,834 and 694,649 individuals respectively from the GIANT study^80,81^, circulating triglyceride levels in up to 1,253,277 European individuals from the Global Lipids Genetics Consortium (GLGC) study^82^, type 2 diabetes (T2D) incidence in up to 2,535,601 multi-ancestry participants from the DIAGRAM study^83^, and whole body fat percentage data (field 23099) in up to 444,608 participants from the UK Biobank^84^. For each GWAS, we identified independent GWAS signals and prioritised causal gene candidates using the “GWAS to Genes” (G2G) framework as recently described^29^ and summarized below.

GWAS summary statistics were filtered to retain common variants with a MAF>0.1%. Quasi-independent genome-wide significant signals were initially selected in 1Mb windows and secondary signals within these loci were then selected via conditional analysis in GCTA v.1.93.2^85^, using an LD reference derived from the UK Biobank study. Primary signals were then supplemented with unlinked (r2<0.05) secondary signals and mapped to proximal NCBI RefSeq genes, within 500kb windows.

Independent signals and closely linked SNPs (r2>0.8) at each GWAS locus were annotated for their closest gene, for coding variants or if they mapped within known enhancers of the identified genes, via activity-by-contact (ABC) enhancer maps^86^. Gene-level associations were tested using Multi-marker Analysis of GenoMic Annotation (MAGMA)^87^ by collapsing all common (MAF>0.1%) coding variants within each gene. Colocalization analyses between GWAS SNPs and expression- or protein-quantitative trait loci (eQTL or pQTL) data were performed via SMR-HEIDI (summary data–based Mendelian randomization-heterogeneity in dependent instruments, v0.68)^88^ and the ABF function within the R package “coloc” v5.1.0^89^. For eQTL analyses, these were applied for specifically enriched tissues (via LDSC-SEG)^90^, as well as cross-tissue meta-analysed GTEx eQTL data v7 (GTEx Consortium, 2015^93^ available via https://gtexportal.org), blood eQTLs^91^ and Brain-eMeta^92^ studies. Lastly, genes at each locus underwent prioritization using the polygenic priority score (PoPS) method^94^. Causal candidate genes were then prioritized by scoring the evidence from all of the above sources. Genes were considered to be confidently implicated by a GWAS signal if supported by at least two of the above sources of evidence (for further details of this G2G (GWAS to genes) approach^29^.

GWAS signals, and any SNPs in high LD (r2>0.8), were mapped (if present) to the human genomic regions syntenic to the mouse DInts regions. Orthologous genes identified as novel regulators of late-stage adipogenesis were also scored using G2G evidence for the same five metabolic GWAS traits.

## Statistical analyses

Statistical analyses were performed as described above, or using GraphPad Prism 10 software. For all tests, P<0.05 was considered significant.

## Data availability

Sequencing data was deposited in NCBI’s Gene Expression Omnibus (GEO) under the accession numbers GEO: GSE234744 (RNA-seq of primary pre-adipocytes and adipocytes isolated from 3 months-old hemizygous Zfp423^GFP^ mice), GSE234747 (PCHi-C-seq of D0 and D7 3T3-L1 cells) and GSE234749 (CAGE-seq of D0 and D7 3T3-L1 cells).

## Acknowledgments

This work was supported by the Medical Research Council (MR/J001562/1 to M.C.; MRC_MC_UU_12014/4 to M.C. and S.E.O.; MRC_MC_UU_12012/5 to the MRC Metabolic Diseases Unit; MRC_MC_UU_12015/2 and MRC_MC_UU_00006/2 to J.R.B.P. and K.K.O.) and the Wellcome Trust (214274/Z/18/Z to S.O.R). This research was also conducted using the UK Biobank Resource under application 9905. S.S. was supported by a UKRI MRC Rutherford Fund Fellowship (MR/T016787/1), a BBSRC Institute Strategic Programme grant (BBS/E/B/000C0421), and a Career Progression Fellowship from the Babraham Institute. F.M. was supported by the Qatar National Research Fund (UREP28-269-1-051 and NPRP14S-0319-210075). A.B. and F.M. were supported by funding from CHLS (College of Health & Life Sciences, Hamad Bin Khalifa University, Doha, Qatar). N.C. was funded by the Frank Edward Elmore Fund, the Association of Physicians of Great Britain & Ireland and the Anatomical Society. A.E., B.S.H. and A.T.C. were funded by MRC Programme Grants MC_UU_12022/1 and MC_UU_12022/8 awarded to A.R.V. The diagram presented in Figure 5A was generated using BioRender (https://www.biorender.com).

## Author contributions

M.C. and S.E.O. designed the project; M.C., S.E.O., F.M., S.O.R., J.R.B.P. and A.R.V. secured the funding; I.S., B.M., A.E., P.G., K.A.K., P.F., S.S., F.M., S.O.R., J.R.B.P., A.R.V., S.E.O. and M.C. designed the experimental setup; I.S., A.E., P.G., A.B., N.C., B.S.H., A.T.C., D.S.F.-T., S.S. and F.M. performed all in vitro and in vivo experimental work; B.M., I.S., K.A.K., L.S., A.E., P.G., A.B., D.S.F.-T., L.V.M., L.S., S.A., D.C., R.S.H., S.W.W., K.K.O., S.S., F.M., A.R.V. and J.R.B.P. performed data analysis and interpretation; I.S., B.M., K.A.K., S.E.O. and M.C. wrote the paper. All other authors discussed the results and edited the manuscript. M.C. managed and supervised all aspects of the study.

