## Supplementary data for "Rewiring of chromatin loops in adipogenesis reveals targets for obesity and diabetes intervention"

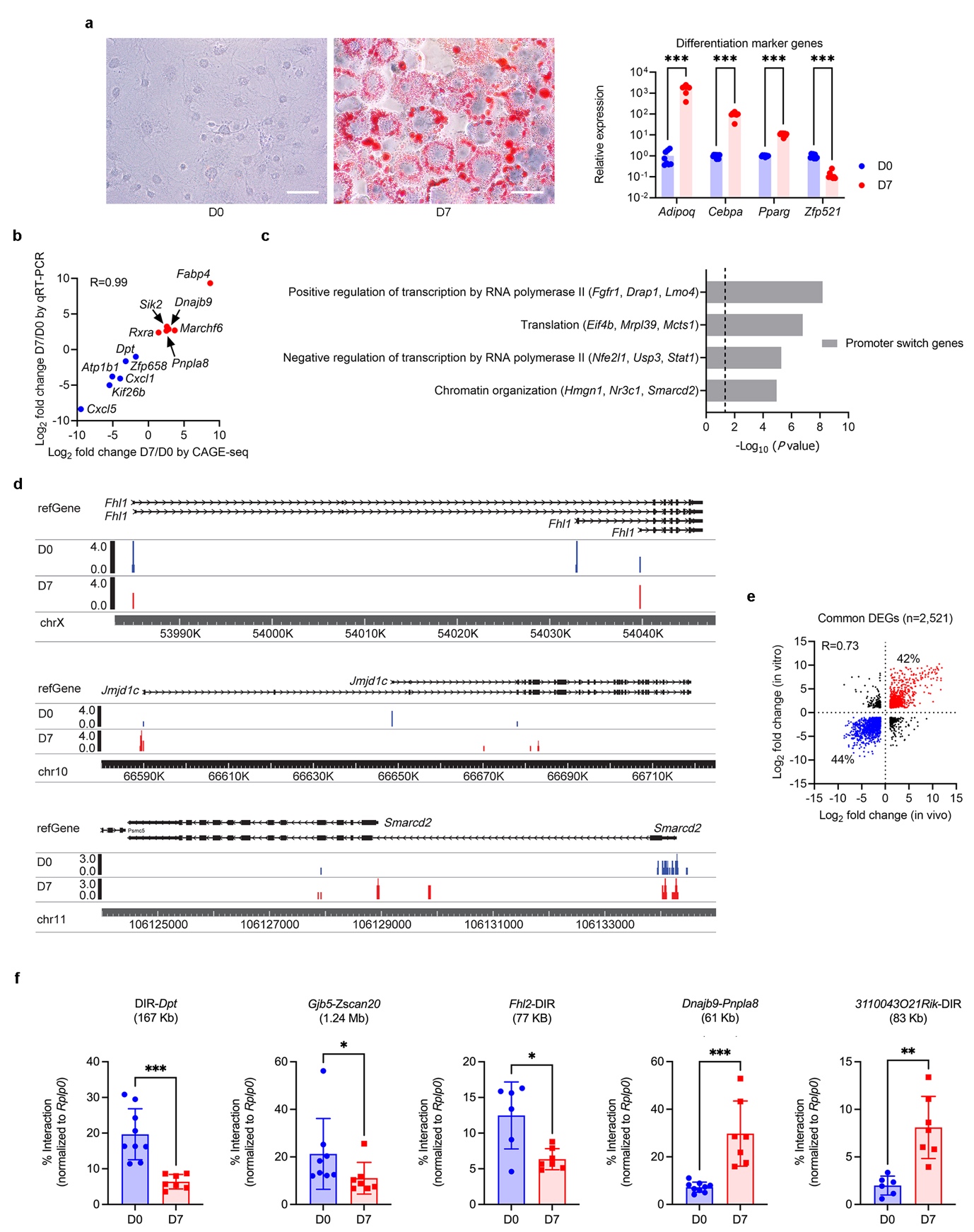

**Extended Data Fig. 1: Transcriptional and epigenetic changes observed in the 3T3-L1 model of in vitro adipogenesis.**

**(a)** Left: Oil Red O staining of 3T3-L1 pre-adipocytes (day 0, D0) and adipocytes (day 7, D7). Scale bars, 20 µm. Right: quantification of mRNA levels of marker genes induced (*Adipoq*, *Cebpa*, *Pparg*) or suppressed (*Zfp521*) during adipogenesis. Error bars represent s.d.; ***P < 0.001 by Mann–Whitney test. **(b)** Linear regression between expression levels measured by CAGE-seq (x axis) and qRT-PCR (y axis) in biological replicates, validating CAGE-seq results. **(c)** DAVID functional annotation of 605 genes showing promoter switching (identified by CAGE-seq) between D0 and D7. The dotted line indicates P = 0.05. **(d)** Examples of genes exhibiting promoter switching (blue, peaks expressed at D0; red, peaks expressed at D7). **(e)** Linear correlation of genes differentially expressed (>2-fold change) in vivo between pre-adipocytes and adipocytes (n = 2,521; x axis) and in vitro between D0 and D7 (y axis). 86% of genes show concordant expression changes (R = 0.73). **(f)** Validation of differential interactions (DInts) between D0 and D7 identified by GOTHiC at five loci, using quantitative 3C (q3C) in biological replicates. Distances between promoters and distal interacting regions (DIR) are indicated in brackets. Error bars represent s.d.; *P < 0.05, **P < 0.01, ***P < 0.001 by Mann–Whitney test.

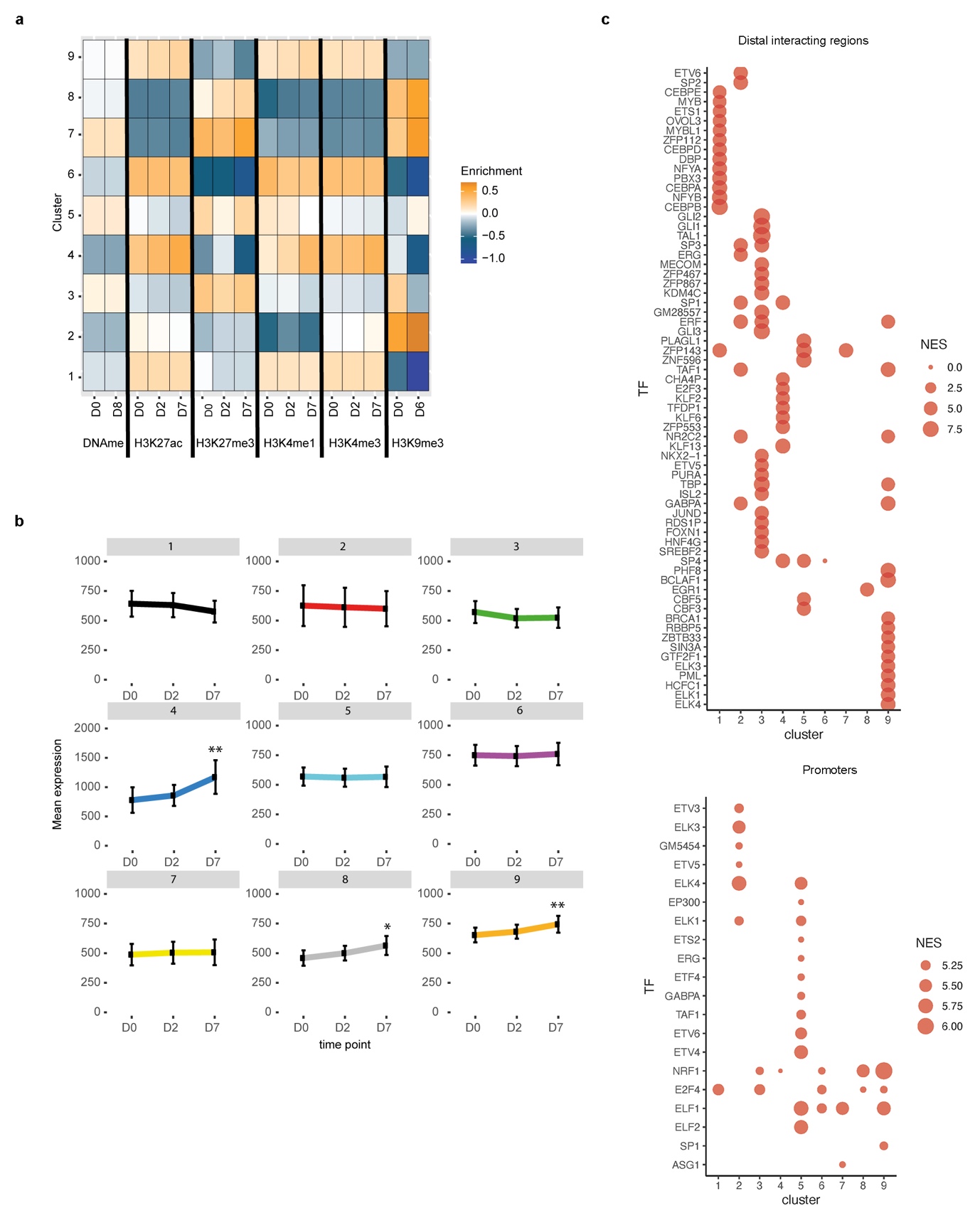

**Extended Data Fig. 2. Supporting data for the cluster analysis of differential promoter-anchored interactions (DInts).**

**(a)** Epigenetic mark enrichment at promoters of genes involved in DInts across differentiation stages (D0, D2, D7) in 3T3-L1 cells, stratified by cluster. Note inter-cluster differences but relative intra-cluster stability per mark during adipocyte differentiation. Enrichments are relative to the average levels across the nine clusters. **(b)** Patterns of gene expression changes per cluster from microarray data during 3T3-L1 differentiation (D0, D2, D7). Data are mean expression levels; error bars indicate 95% confidence intervals. *P < 0.05, **P < 0.01 by one-way ANOVA with Tukey’s multiple comparisons test. **(c)** Dot plots showing transcription factor binding site enrichment at distal interacting regions (top) or promoters (bottom) of genes within specific DInt clusters. NES, normalized enrichment scores.

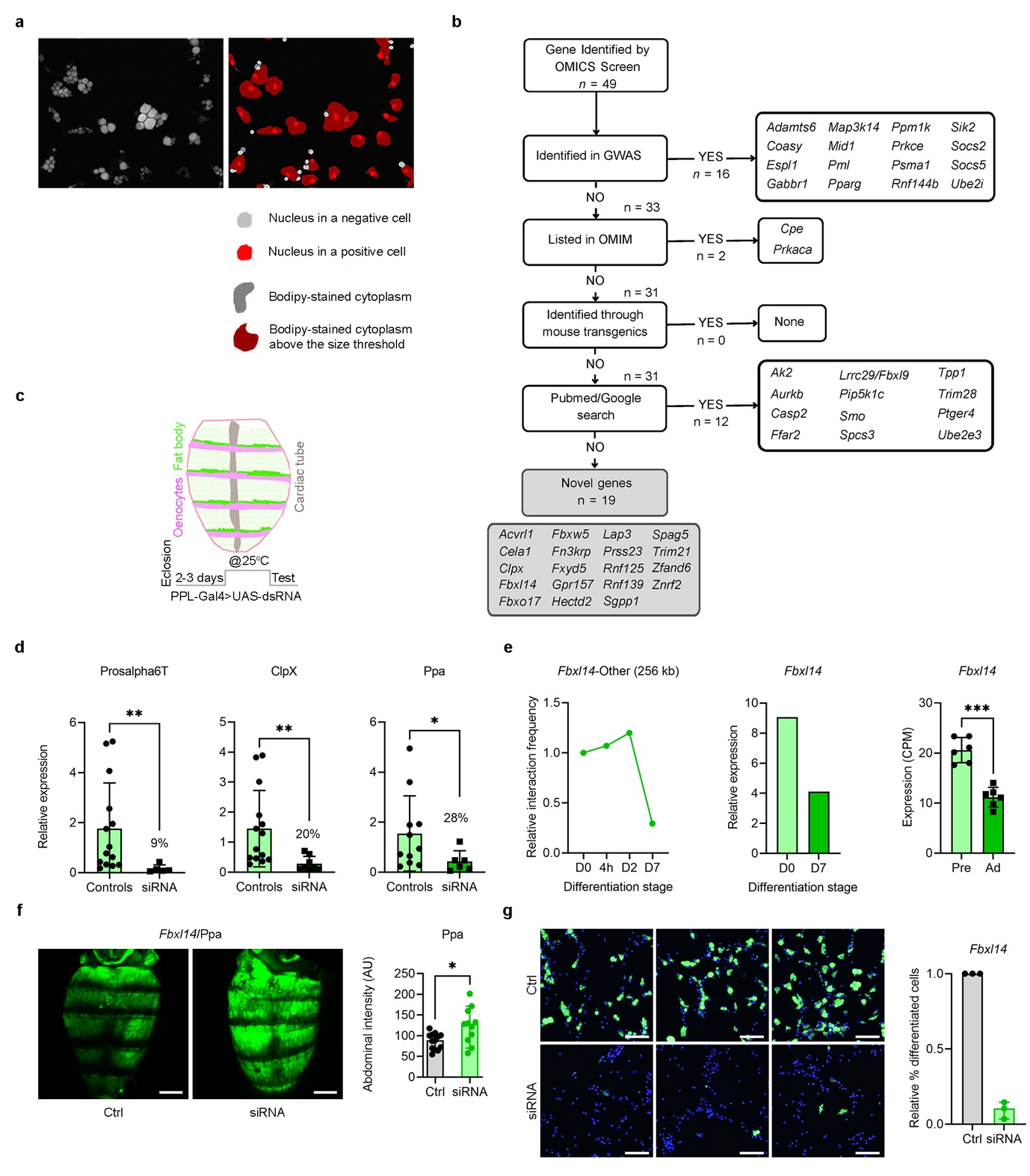

**Extended Data Fig. 3. Supporting data for the identification and validation of novel regulators of late-stage adipogenesis.**

**(a)** Assessment of lipid accumulation upon siRNA knockdowns in vitro in OP9-K cells. Left: Bodipy-stained OP9-K cells. Right: image overlay highlighting nuclei and Bodipy-stained cytoplasm (see Methods). **(b)** Flowchart illustrating manual curation steps and decision points for identifying novel regulators of late-stage adipogenesis. **(c)** Assessment of lipid accumulation upon siRNA knockdowns in vivo in Drosophila fat body. Top: schematic of Drosophila anatomy showing fat body location. Bottom: outline of experimental setup (see Methods). **(d)** Efficiency of siRNA knockdowns in Drosophila for *Prosalpha6T*, *ClpX* and *Ppa*, measured by qRT-PCR in whole flies; values in parentheses indicate expression reduction relative to controls. mRNA normalized to *Rpl32*. Error bars, s.d.; *P < 0.05, **P < 0.01 by Mann–Whitney test. **(e)** Quantification of Fbxl14 promoter-anchored chromatin loop by GOTHiC during 3T3-L1 adipogenesis (D0, 4h, D2, D7; left), Fbxl14 expression by CAGE-seq in pre-adipocytes (D0) and adipocytes (D7; middle), and Fbxl14 expression by RNA-seq in primary murine pre-adipocytes and adipocytes (right). Error bars, s.d.; ***P < 0.001. **(f)** Left: representative images of Bodipy-stained Drosophila abdomens following siRNA treatment (scale bars, 200 µm). Right: quantification of Bodipy intensity in fat body. Error bars, s.d.; **P < 0.01, ***P < 0.001 by unpaired t test with Welch’s correction. **(g)** Left: representative images of OP9-K adipocytes stained with Bodipy (green) and DAPI (blue) following siRNA knockdown of *Fbxl14* (scale bars, 50 µm). Right: quantification of percentage differentiated cells relative to controls (Ctrl), normalized to 1. Error bars, s.d.

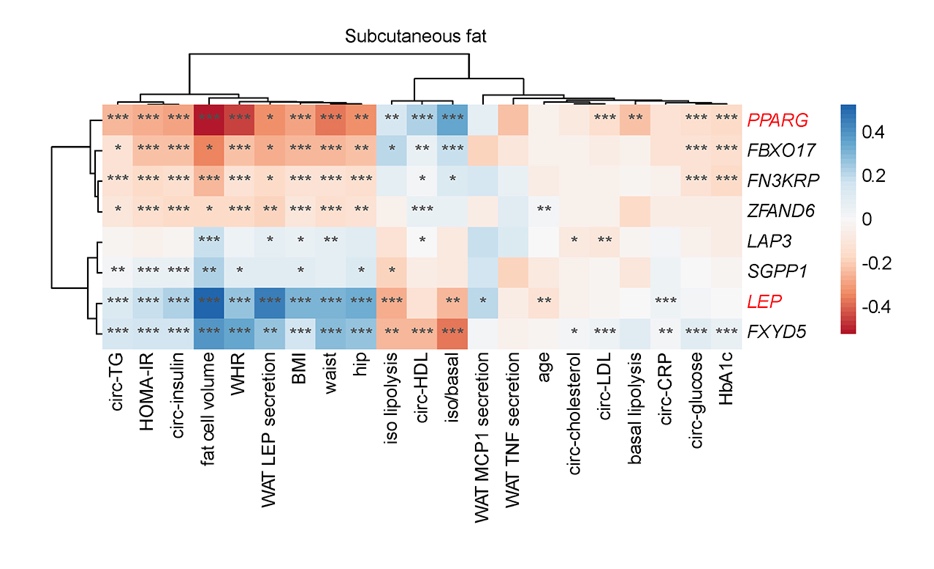

**Extended Data Fig. 4. Interrogation of the Adipose Tissue Knowledge Portal for evidence supporting the involvement of six candidate genes as novel regulators of late-stage adipogenesis in human.**

The heatmap shows Pearson correlation analyses between six novel regulators of late-stage adipogenesis, plus *PPARG* and *LEP* (in red, as controls), and relevant clinical parameters from the “clinical module” of the Adipose Tissue Knowledge Portal. Colour intensity indicates correlation strength; stars (*, **, ***) denote statistical significance levels. Clustering was performed using the ward.D method.

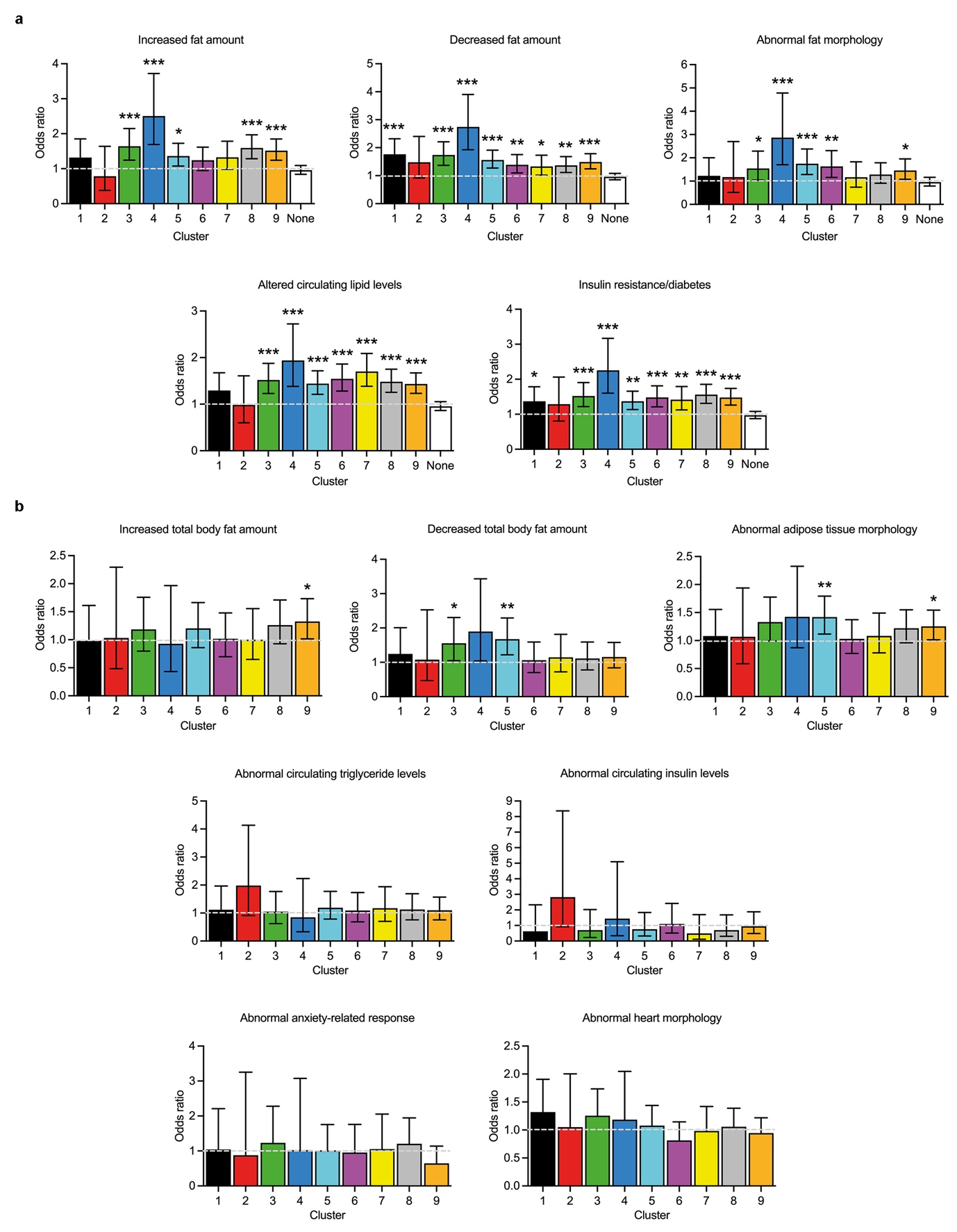

**Extended Data Fig. 5. Genes engaged in DInts during 3T3-L1 adipogenesis are enriched in obesity and diabetes phenotypes in mouse.**

**(a)** MGI (Mouse Genome Informatics) data showing significant enrichment of genes per nine DInt clusters whose loss- or gain-of-function mutations cause obesity- or diabetes-related phenotypes in vivo. The “None” column includes phenotyped genes not belonging to any DInt cluster, serving as a specificity control. **(b)** IMPC (International Mouse Phenotyping Consortium) data showing enrichment of genes per DInt cluster whose knockout leads to obesity- or diabetes-related phenotypes in vivo. The last two panels (abnormal anxiety-related response and abnormal heart morphology) serve as negative controls for phenotype specificity. Columns indicate odds ratios; error bars show 95% confidence intervals. *P < 0.05, **P < 0.01, ***P < 0.001 by Fisher’s exact test. Dotted lines indicate odds ratio = 1.

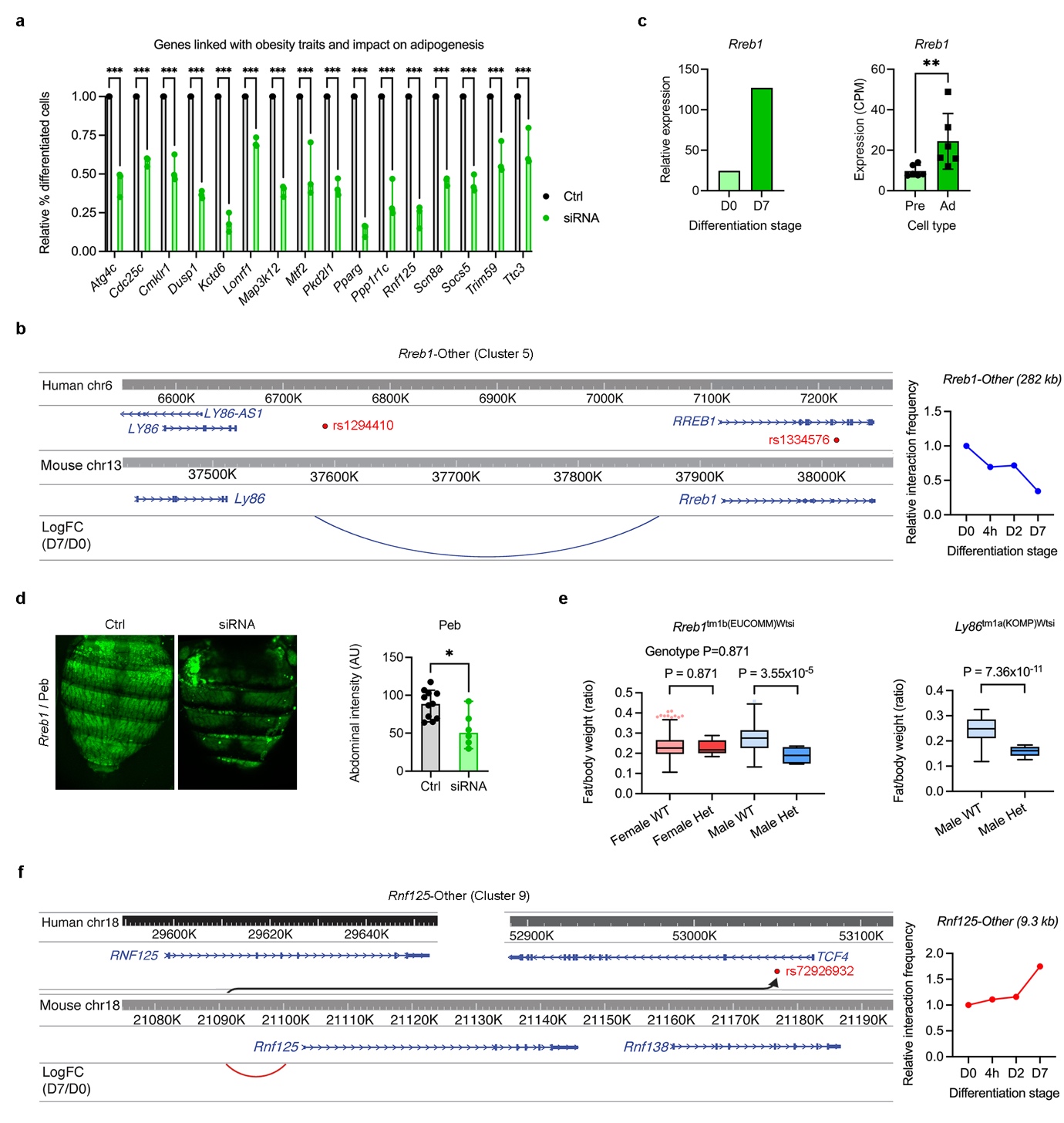

**Extended Data Fig. 6. Additional validation of novel regulators of late adipogenesis using siRNA knockdowns and analyses of IMPC datasets.**

**(a)** Genes associated with obesity and T2D in human for which siRNA knockdown in mouse OP9-K cells leads to reduced lipid droplet formation during adipocyte differentiation. Data are shown as % differentiated cells relative to controls (Ctrl), normalized to 1. Error bars, s.d.; ***P < 0.001. **(b)** *Rreb1* upregulation during in vitro adipogenesis in 3T3-L1 cells quantified by CAGE-seq (left) and during in vivo adipogenesis measured by RNA-seq (right). Error bars, s.d.; **P < 0.01. **(c)** Left: synteny analysis between a region on human chromosome 6 (containing rs1294410 linked to WHRadjBMI by GWAS implicating *LY86* as causative gene) and homologous region on mouse chromosome 13 (showing cluster 5 DInt linking *Rreb1* promoter to region homologous to rs1294410). *RREB1* is also independently associated with WHRadjBMI via rs1334576 in GWAS Catalog. Right: dynamics of cluster 5 DInt *Rreb1*-Other during 3T3-L1 adipocyte differentiation quantified by GOTHiC. **(d)** siRNA knockdown of *Peb* (Drosophila homologue of *Rreb1*) reduces lipid accumulation in fat body. Left: representative Bodipy-stained Drosophila abdomens post-siRNA treatment (scale bars, 200 µm). Right: quantification of Bodipy intensity in fat body. Error bars, s.d.; *P < 0.05 by unpaired t test with Welch’s correction. **(e)** Left: IMPC data showing significant reduction of fat content in heterozygous males carrying mutant *Rreb1* allele (*Rreb1^tm1b(EUCOMM)Wtsi*). Right: IMPC data showing significant reduction of fat/body weight ratio in heterozygous males carrying mutant *Ly86* allele (*Ly86^tm1a(KOMP)Wtsi*). **(f)** Left: synteny analysis between human chromosome 18 region (rs72926932 linked to T2D by GWAS implicating *TCF4*) and homologous mouse chromosome 18 region (showing cluster 9 DInt linking *Rnf125* promoter with region homologous to rs72926932 inside an intron of *TCF4*). Right: dynamics of cluster 9 DInt *Rnf125*-Other during 3T3-L1 adipocyte differentiation quantified by GOTHiC. Curved arrow indicates positional change of distal interacting fragment from upstream *Rnf125* in mouse to intron of *TCF4* in human.

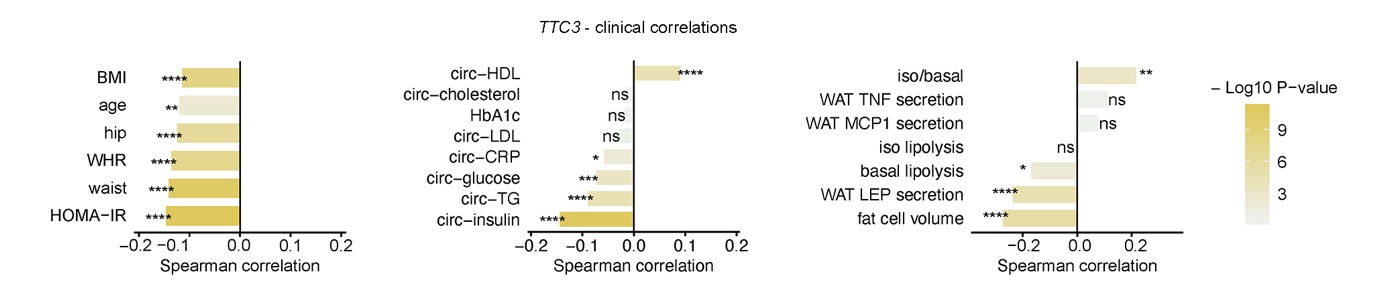

**Extended Data Fig. 7. Interrogation of the Adipose Tissue Knowledge Portal for evidence supporting the involvement of TTC3 as a novel regulator of late-stage adipogenesis in humans.**

The graphs depict correlation analyses between *TTC3* and relevant clinical parameters, obtained using the “clinical module” of the Adipose Tissue Knowledge Portal. Colour intensity reflects the strength of Spearman correlations. Asterisks denote levels of statistical significance (*P < 0.05, **P < 0.01, ***P < 0.001; ns, not significant).

**Supplementary Table 1. Identification of differentially expressed genes during 3T3-L1 adipogenesis by CAGE-seq and genes that associate a promoter switch.**

**Supplementary Table 2. Identification of 9 clusters of DInts in 3T3-L1 cells and analyses of transcription factor binding and gene ontology performed for each cluster.**

**Supplementary Table 3. siRNA screen of 2,905 “druggable” genes for impact on lipid accumulation in OP9 cells and manual curation for the identification of 19 novel regulators of late-stage adipogenesis.**

**Supplementary Table 4. GWAS for the 19 novel regulators of late-stage adipogenesis.**

**Supplementary Table 5. DInt genes associated with obesity and diabetes phenotypes based on MGI and IMPC data.**

**Supplementary Table 6. Multi-trait “GWAS to Genes” (G2G) analysis on human chromosomal regions with preserved synteny of homologous chromosomal fragments implicated in DInts in the mouse.**

**Supplementary Table 7. Primers used for qRT-PCR or 3C assays.**

| Primers used for qRT-PCR in 3T3-L1 cells | | | | | |
| --- | --- | --- | --- | --- | --- |
| Gene | Primer | Sequence (5’-3’) | Primer | Sequence (5’-3’) | Amplicon (bp) |
| *Adipoq* | F | GCCACTTTCTCCTCATTTCTGT | R | CTTGCAACAGTAGCATCCTGAG | 82 |
| *Atp1b1* | F | GGAAGGCAGCTGGAAGAAATT | R | CAGCCAGGCAGCCATAAAATA | 119 |
| *Cebpa* | F | ATAGACATCAGCGCCTACATCG | R | CTCCCGGGTAGTCAAAGTCAC | 142 |
| *Cxcl1* | F | CCACACTCAAGAATGGTCGC | R | GCAGTCTGTCTTCTTTCTCCG | 116 |
| *Cxcl5* | F | GAAGGAGGTCTGTCTGGATCC | R | TTCTATTGAACACTGGCCGTT | 129 |
| *Dnajb9* | F | TAGCCATGAAGTACCACCCTG | R | AGCACTGTGTCCAATTGTGTC | 134 |
| *Dpt* | F | TACCAGAAGTGCTCCAACAATG | R | GCTTGGGTACTCTGTTGTCATC | 147 |
| *eRNA (Fabp12)* | F | GTGCCTTAGAGAGTTCATGCAC | R | CCAGTAAGTGCGTTGAGGTACA | 89 |
| *Fabp4* | F | AGCTTGTCTCCAGTGAAAACT | R | CCCCATTTACGCTGATGATCA | 114 |
| *Kif26b* | F | AGAGCCACCATGAATTCGGTA | R | GTAAGCCTTGCGGTACCAGC | 144 |
| *Marchf6* | F | CAGACAAGATGGACACCGCC | R | ACTGCCAGTACATACACAAGGA | 101 |
| *Pnpla8* | F | GCCTTAAGAAGAACAGCCGAC | R | TCCTTAACTTGTCTCAGCCGT | 146 |
| *Pparg* | F | ACCTGAAGCTCCAAGAATACCA | R | CCTGTTGTAGAGCTGGGTCTTT | 89 |
| *Ppia* | F | CTGAGCACTGGAGAGAAAGGAT | R | TTATGGCGTGTAAAGTCACCAC | 98 |
| *Rplp0* | F | TGAGTACACCTTCCCACTTACTGA | R | CTCTTCCTTTGCTTCAGCTTTG | 145 |
| *Rxra* | F | GGGCATGAGTTAGTCGCAGA | R | GAGAGTTGAGGGACGAAGAGT | 88 |
| *Sik2* | F | CAACTTTGCCGTGGTGAAGC | R | GTGAGGATGGTCGAGCATTTT | 148 |
| *Zfp521* | F | GTGTCACCTGATAGAGCACAGC | R | TGCTGTTGCAACTTGTTAGCTT | 99 |
| *Zfp658* | F | CGTCCTGTGCAGAGTTCTCT | R | GAACTGGAAGCTGGTTGAGTG | 149 |
| Primers used for verifying efficiency of *Hind*III digestion and DNA ligation | | | | | |
| Locus | Primer | Sequence (5’-3’) | Primer | Sequence (5’-3’) | Amplicon (bp) |
| q*Calr*-*Hind*III | F | ACTTACACCAACTCTTAAGGTACC | R | ATGAACTGCCCTATCCTGAGTC | 126 |
| q*Calr*-Int | F | CTCCTGCCCATTCTTCAACCTA | R | GTTGGACACTTAGCTAGCATGG | 123 |
| *Calr* | F | TCATGAGTTCCCCACATCTTTG | R | CTGTGGGCACCAGATGTGTAAAT | 257 |
| *Gapdh* | F | TATCAAGGGTGCCCGTCACCTTCAGC | R | GGGCTTTTATAGCACGGTTATAAAGT | 305 |
| *Hist1h2ae- Hist1h3e* | F | GGGTAATGGTGTCACTAACTGG | R | GGGTTTGATGAGTTGGTGAAG | 176 |
| *Hoxa7-Hoxa13* | F | GGAGGAGGGAAAAGGAGTGATT | R | CAGGCATTATTTGCTGAGAACG | 400 |
| *Hist1h2ae- Hist1h4i* | F | GGGTAATGGTGTCACTAACTGG | R | TTGGGCCAAAGCCTATATGA | 161 |
| Primers used for q3C assays in 3T3-L1 cells | | | | | |
| Interaction | Primer | Sequence (5’-3’) | Primer | Sequence (5’-3’) | Amplicon (bp) |
| *3110043O21Rik*-DIR | F | CTACCATACCCTGCTGCTGT | R | GGATCTCCCTATGTAGTCCAAC | 113 |
| *Dnajb9-Pnpla8* | F | AATTCAAAACTGCACGCCCT | R | GGAGCACTTCTTCGGTAAACC | 150 |
| *Fabp4*-DIR | F | CTTTGCTACAACGGGAACATG | R | CATCCACTCAACAAAACCCCT | 150 |
| *Fhl2*-DIR | F | TGTGTGTAGCCAATTTTGCAGT | R | CCACTGAGAGCAACTGGATTTT | 141 |
| *Gjb5-Zscan20* | F | GCTATACAGAGGACAGAGGACC | R | CGTCTCAGAAGGAAGTGGTTTT | 145 |
| DIR-*Dpt* | F | GGGATGGTTTTCTTCTGAGCA | R | TGAGTCACAAGCTGGGTTTTG | 114 |
| *Rplp0* | F | TCTCTTCCCTGATCAGAGTCAC | R | TCCCTAGAGTTGTGTTATTTAGTAGT | 121 |
| Primers used for qRT-PCR in *Drosophila* | | | | | |
| Gene | Primer | Sequence (5’-3’) | Primer | Sequence (5’-3’) | Amplicon (bp) |
| Clpx | F | CTAACGCACACTCAAGAACGC | R | GGCTCTTCGCTCCACTACC | 96 |
| Ppa | F | CCAGTGAGGGATTTGGGTCG | R | CAGGCAGTTGAACAGGCTC | 135 |
| Prosalpha6T | F | AATGGAGGCGGTGAAACAGG | R | CGTGTTCGTGTCCTTTGAGGT | 100 |
| Rpl32 | F | CGGATCGATATGCTAAGCTGT | R | GCGCTTGTTCGATCCGTA | 184 |
